# Microevolution in the major outer membrane protein OmpA of *Acinetobacter baumannii*

**DOI:** 10.1101/711606

**Authors:** Alejandro M. Viale, Benjamin A. Evans

## Abstract

*Acinetobacter baumannii* is nowadays a relevant nosocomial pathogen characterized by multidrug resistance (MDR) and concomitant difficulties to treat infections. OmpA is the most abundant *A. baumannii* outer membrane (OM) protein, and is involved in virulence, host cell recognition, biofilm formation, regulation of OM stability, permeability, and antibiotic resistance. OmpA members are two-domain proteins with an N-terminal eight-stranded β-barrel domain with four external loops (ELs) interacting with the environment, and a C-terminal periplasmic domain binding non-covalently to the peptidoglycan. Here, we combined data from genome sequencing, phylogenetic, and multilocus sequence analyses from 242 strains of the *Acinetobacter calcoaceticus/Acinetobacter baumannii* complex (ACB), 222 from *A. baumannii*, to explore *ompA* microevolutionary divergence. Five major *ompA* variant groups were identified (V1 to V5) comprising 50 different alleles coding for 29 different proteins. Polymorphisms were concentrated in 5 regions corresponding to the four ELs and the C-terminal end, and provided evidence for different intra-genic recombination events. *ompA* variants were not randomly distributed across the *A. baumannii* phylogeny, with the most frequent V1a1 allele almost exclusive to clonal complex 2 (CC2) strains and the second most frequent V2a1 allele found in the majority of CC1 strains. Evidence was found for assortative exchanges of *ompA* alleles not only between different *A. baumannii* clonal lineages, but also different ACB species. Within *A. baumannii ompA* non-synonymous substitutions were concentrated in the ELs regions, but were more abundant in the transmembrane regions between different *Acinetobacter* species. The overall results have implications for *A. baumannii* evolution, epidemiology, virulence, and vaccine design.

**Importance:** *Acinetobacter baumannii* is an increasing MDR threat in nosocomial settings associated with prolonged hospitalization and concomitantly increased healthcare costs. The main *A. baumannii* OM protein, OmpA, is a multifaceted two-domain protein implicated in host cell recognition and adhesion, cytotoxicity, biofilm formation, and as a slow porin for antibiotics and small hydrophilic nutrients. *A. baumannii* OmpA has been proposed as a potential target for anti-virulence drugs and as a vaccine candidate. Given the many interactions of this protein with environmental factors including host defenses, it is certainly subjected to many selective pressures. Here, we analyzed the microevolution of this OM protein in the *A. baumannii* population to obtain clues on the extent to which selection in the clinical setting has shaped this protein. The results provide relevant information on the main causes driving evolution of this protein, with potential implications in *A. baumannii* epidemiology, virulence, and vaccine design.

## Introduction

The genus *Acinetobacter* (family *Moraxellaceae*, class *Gammaproteobacteria*) is composed of Gram-negative bacterial organisms of ubiquitous environmental distribution (1). It includes a number of phylogenetically closely-related species generally associated with the human environment grouped into the so-called *Acinetobacter calcoaceticus/Acinetobacter baumannii* complex (ACB). Of these, *A. baumannii* represents the most problematic nosocomial species generally affecting critically-ill patients (1–6). Molecular typing studies have indicated that infective *A. baumannii* strains belong to a limited number of globally-distributed clonal groups/complexes (CCs) of recent emergence (2–9). The reasons for the success of CC strains as human pathogens are still poorly understood, but are thought to include the general resilience of the species, an increased ability to colonize the human environment, and their propensity to acquire exogenous genetic material (1–9). Thus, CC strains not only show multi-drug resistance (MDR) phenotypes from the acquisition of different mobile genetic elements enriched in resistance genes, but can also rapidly modify virulence-related surface structures recognized by host defenses as the result of recombination events involving discrete genomic loci (2–6, 9–12). This has left very few treatment options available (2–6, 10, 13) and has led to the alternative search of targets for anti-virulence drugs or vaccines, and in this context a potential candidate is represented by the major *A. baumannii* outer membrane (OM) protein, OmpA (6, 14–22).

*A. baumannii* OmpA belongs to the widely distributed OmpA/OprF family of OM proteins from which *Escherichia coli* OmpA and *Pseudomonas* OprF are well-characterized examples (23, 24). These proteins are structured in two domains: an N-terminal eight-stranded β-barrel embedded in the OM exposing four loops to the environment (ELs), and a C-terminal periplasmic domain that binds the peptidoglycan and contributes to cell envelope stability (23–25). *E. coli* OmpA ELs play important roles in target cell recognition and bacterial survival once internalized, and variations in EL2 and EL3 composition or length affects the invasive potential of the bacterial cells (23). Given the many interactions of ELs with environmental factors, it is not surprising that they represent the less conserved protein regions with both mutation and recombination providing for the observed variabilities (23, 26, 27).

*A. baumannii* OmpA has been implicated in a multitude of functions that range from structural roles, a slow porin for small hydrophilic nutrients and antimicrobials, and virulence including fibronectin binding, biofilm formation, adhesion to host cells, and cytotoxicity (3, 6, 18, 19, 24, 28–36). It has also been shown to interact with host defenses, inducing innate immune responses, the production of antibodies both in *A. baumannii*-infected patients and experimentally-infected mice, and to bind antimicrobial peptides via the ELs (14–22, 37). It follows that there are different selective pressures including trade-off mechanisms (23–27, 38) acting on *A. baumannii* OmpA. However, to our knowledge no systematic studies have been conducted on *A. baumannii* OmpA microevolution. Here, we analyzed the polymorphism of *A. baumannii* OmpA and the roles of mutation and recombination in defining this variability, to analyze the extent to which selection in the clinical setting has shaped this protein.

## Results

### OmpA polymorphism in the A. baumannii population

*ompA* sequences were extracted from 242 genomes of *Acinetobacter* species deposited in databases, of which 222 were *A. baumannii* (Table S1 and Table S2). A maximum likelihood phylogeny based upon the alignments of the corresponding translated sequences identified five well-defined OmpA variant groups (bootstrap support ≥93 % in all cases), hereafter called V1 to V5 (Figure 1). In *A. baumannii*, each of the identified variant groups was composed of a different number of alleles sharing high levels of amino acid sequence identity (>97%) between them (Table S2). Between these different variants the protein sequence identities ranged from ca. 94 % for V1 and V2 to ca. 85 % between them and the most divergent group V3 (Table S3). An unequal distribution of *ompA* variants was found, with almost 89 % of the *A. baumannii* strains carrying alleles from only V1 and V2 (Table 1, Table S1). Also, *A. baumannii* V1 and V4 variants each encompassed two sub-variant groups designated V1_1 and V4_1, respectively (Table 1, Table S1, Table S2, Figure 2, Figures S1 and S2). Each subgroup could be distinguished by the presence/absence of a 6 amino acid stretch (AAAPAA) close to the C-terminal end, thus defining QEAAAPAAAQ or QQAQ as alternate C-terminal sequences in the corresponding proteins. The origins of this stretch will be discussed in detail below.

**Figure 1:**
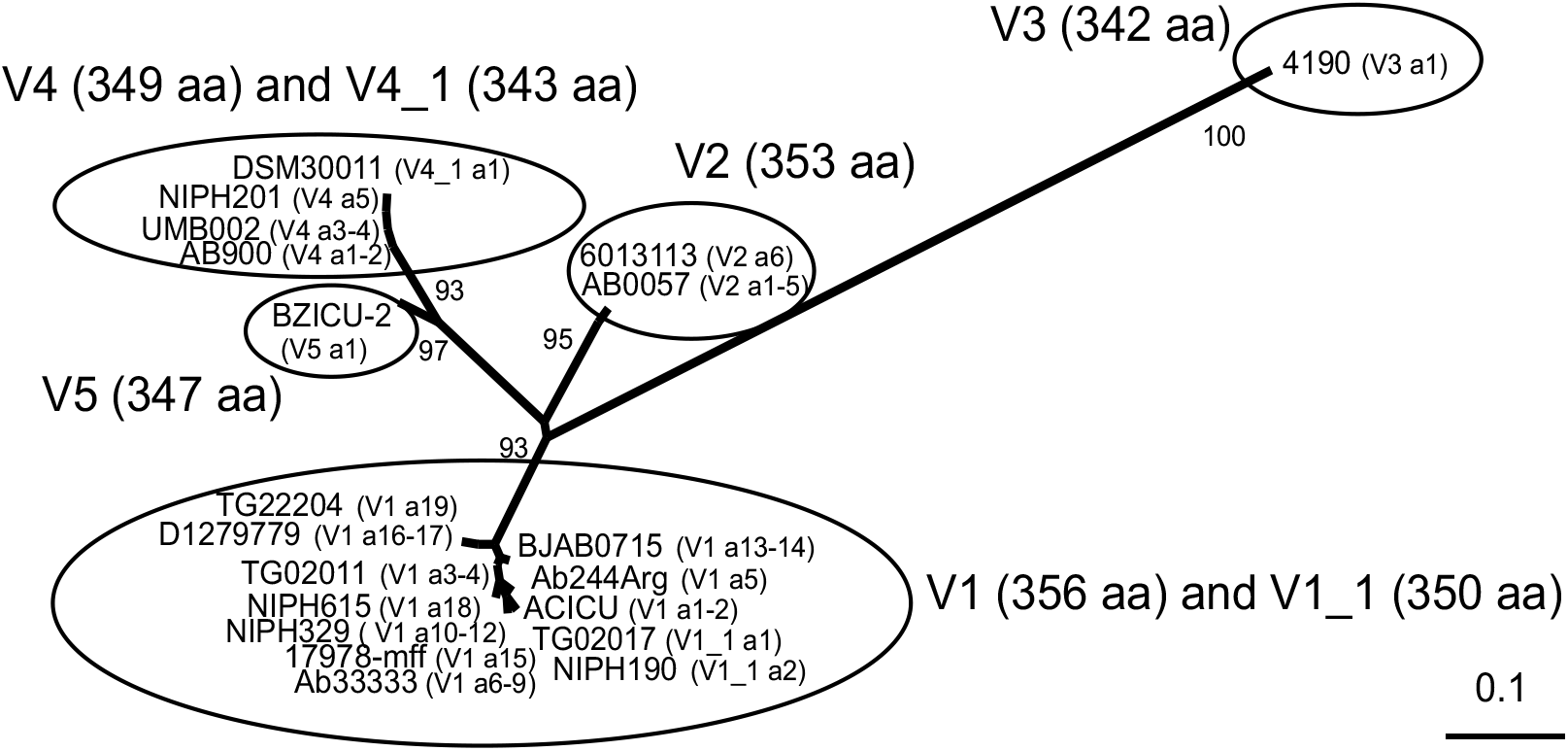
Co-existence of different OmpA variants and sub-variants in the *A. baumannii* population. An unrooted maximum-likelihood (ML) phylogenetic tree was constructed using PhyML (http://phylogeny.lirmm.fr/phylo_cgi/index.cgi) using alignments of translated *ompA* sequences of the 21 indicated *A. baumannii* isolates representing all OmpA protein alleles identified in this species. The lengths of the branches are proportional to the evolutionary distance, with the scale bar (estimated changes per site) shown at the bottom right. Bootstrap support (percentages of 100 resamplings) for the different clusters are indicated at the corresponding branches. The analysis shows that OmpA proteins comprise 5 well-defined similarity groups of alleles (V1 – V5, indicated by ovals) each defining a particular variant. In addition, V1 and V4 encompass sub-variant groups (designated V1_1 and V4_1, respectively) differing from the corresponding V proteins at their C-terminal motifs. The length in amino acid residues of the corresponding variant and sub-variant proteins is indicated for each cluster.

**Table 1:**
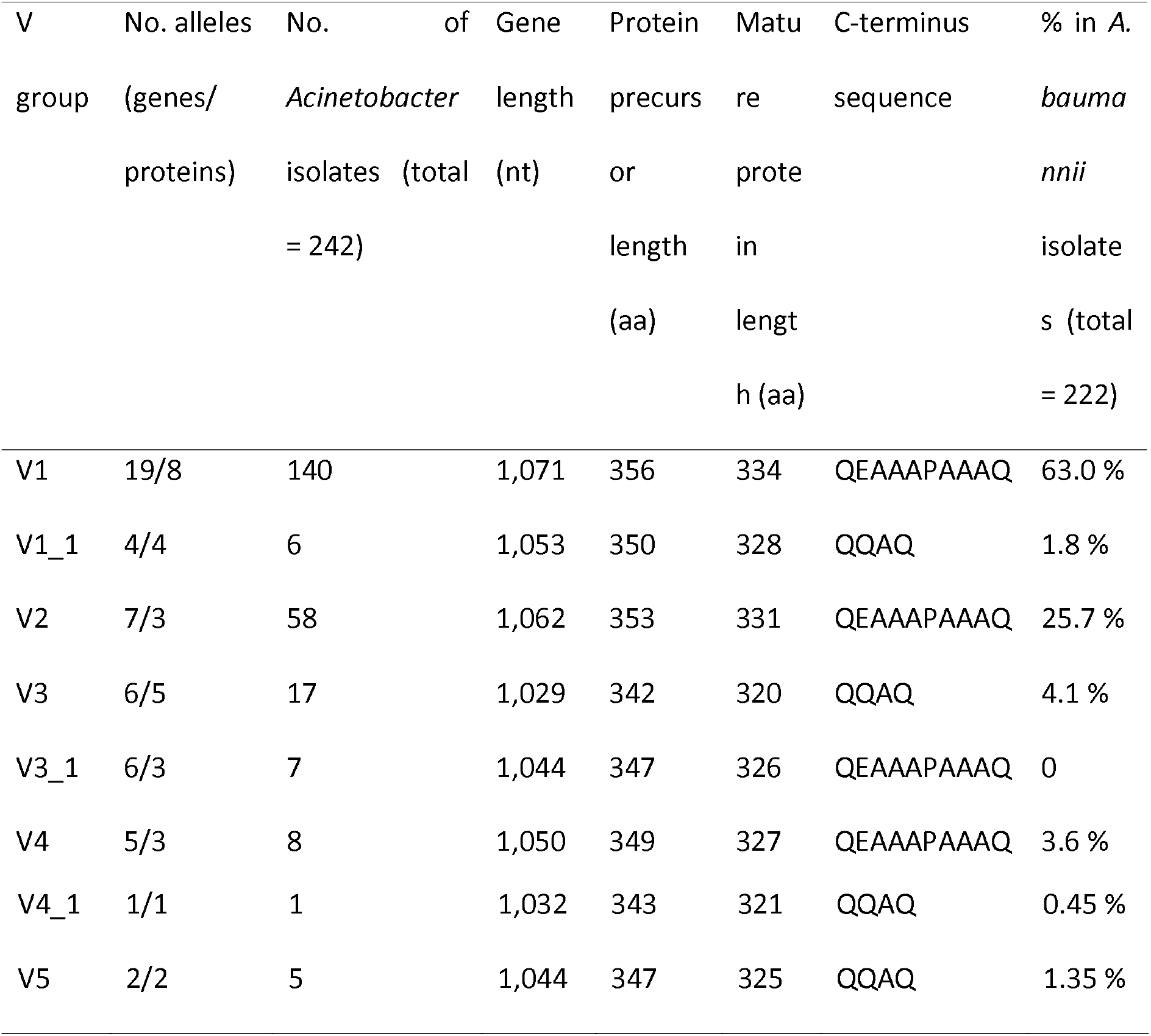
Characteristics of the different OmpA variant (V) groups.

The OmpA V1 group was by far the most frequently observed (63 %) among the *A. baumannii* population analyzed (Table 1). It encompassed 19 gene alleles differing by a maximum of 18 nucleotides, translated to 8 different OmpA proteins differing from 1 to 4 amino acids. Of note, the *A. baumannii* ATCC type strains 19606 and 17978, isolated from human clinical materials around 1950 (39, 40), display the V1 alleles V1a8 and V1a15, respectively (Table S1). This dates the association of V1 *ompA* with *A. baumannii* clinical strains prior to the emergence of the main epidemic clones in the 1960s (5, 9). A sub-variant of V1, designated V1_1 whose alleles display QAAQ at their C-terminal end, were found among a small percentage (1.8 %) of *A. baumannii* strains (Table 1). Two more divergent V1_1 alleles, as judged by the number of amino acid substitutions as compared to V1a1, were also identified in the non-*baumannii* ACB strains *A. nosocomialis* OIFC021 and *A. seifertii* GG2 (Table S2).

V2 was the second most frequent OmpA variant found in 25.7 % of the *A. baumannii* strains analyzed, with 6 different alleles in *A. baumannii* (Table 1 and Table S1). A seventh and more divergent allele (V2a7) was located in *A. calcoaceticus* ANC3811 (Table S2).

The OmpA V3 group had a much lower representation (4.1 %) in *A. baumannii* (Table 1), with all these strains showing V3a1 displaying the “short” QQAQ C-terminal sequence (Figures S1 and S2). Five other V3 alleles were detected among various non-*baumannii* ACB species (Table S2). Notably, a subgroup of V3 alleles (designated V3_1) was also found among *A. pittii, A. calcoaceticus*, and *A. oleivorans* strains (Table S2).

The OmpA V4 group had a low representation in *A. baumannii* (3.6 %, Table 1 and Table S1). It was composed of five alleles displaying the “long” QEAAAPAAAQ C-terminus differing by a maximum of 14 nucleotides translated to only 3 different proteins (Table S2). A V4_1 sub-variant displaying the shorter QQAQ C-terminal end was found in the *A. baumannii* DSM30011 strain (Table 1 and Table S2). DSM30011 was isolated before 1944 from a desert plant source and, in agreement with an environmental origin, is susceptible to most clinically-employed antimicrobials including folate synthesis inhibitors (41).

Finally, V5 had a very low representation in *A. baumannii* (1.35 %, only 3 strains all showing the same V5a1 allele, Table 1). A second V5 allele (V5a2) was found in an *A. pittii* strain (Table S2).

### A. baumannii OmpA polymorphism

A Shannon entropy plot of all aligned *A. baumannii* OmpA variant alleles (Figure 2A) showed that approximately 60 from a total of almost 357 amino acid positions (17 %) are polymorphic. This polymorphism was concentrated in five well-defined regions, four of them overlapping with the four extracellular loops located at the N-terminal domain (EL1 – EL4) and the fifth at the C-terminal end (Figure 2A). Amino acid (Figure 2A) and nucleotide (Figure 2B) alignments show that the differences in length among variants are due to indels located in EL1, EL2, and EL3, and in the C-terminal end.

**Figure 2:**
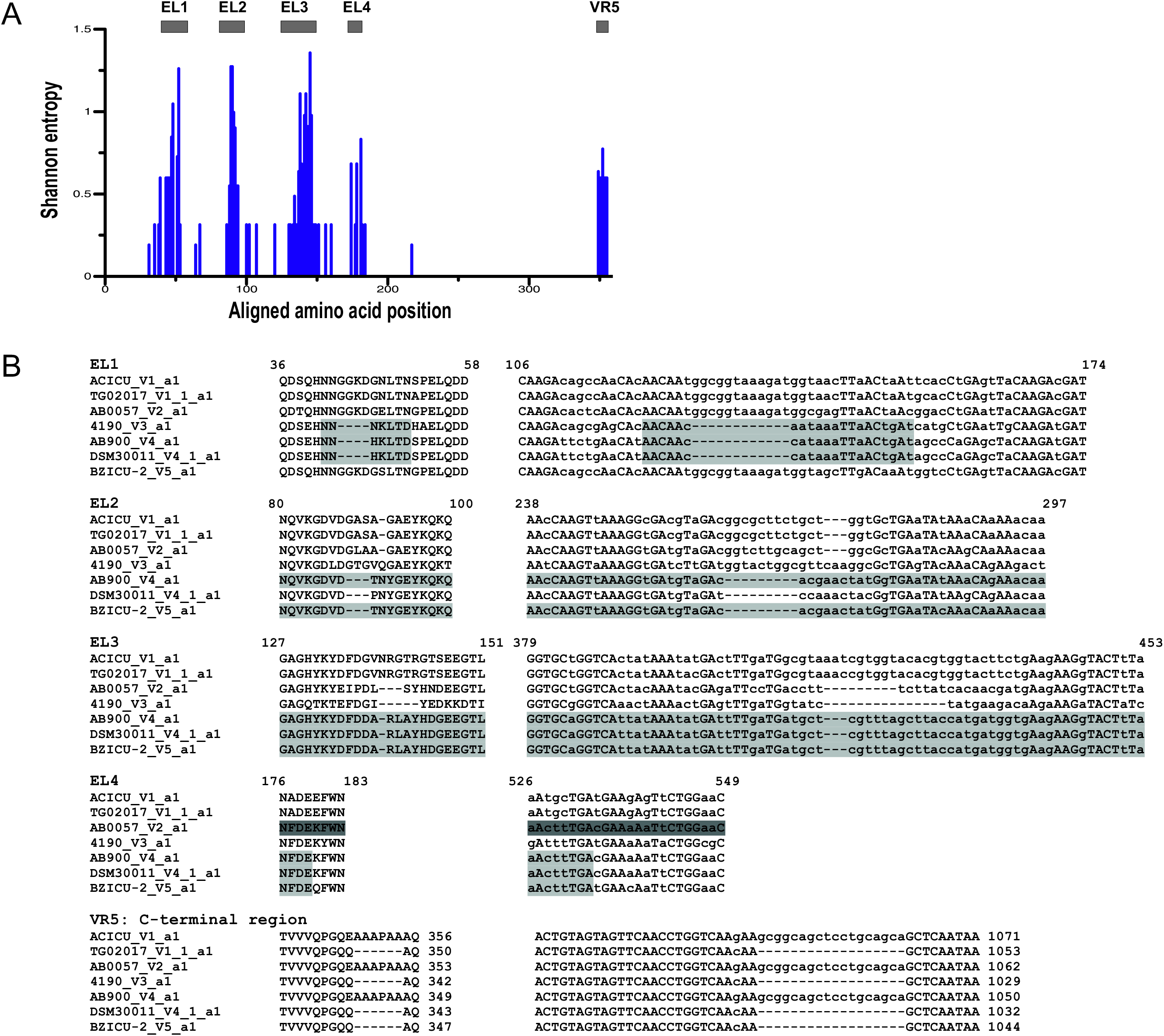
Sequence variability between *A. baumannii* OmpA variants and sub-variants. (A) Shannon entropy variation along the amino acid alignments of the OmpA variant and sub-variant representative alleles shown in Figure 1. The program available at http://www.hiv.lanl.gov/content/sequence/ENTROPY/entropy.html was used for entropy computations. The heights of the bars are proportional to the degree of amino acid variation at a particular location in the alignments. The span of the predicted external loops EL1 to EL4 at the N-terminal domain and the variable C-terminal motif (VR5) are indicated by closed bars above the figure. Alignments are numbered from the corresponding N-terminal regions including the signal peptide. (B) Amino acid (left columns) and corresponding nucleotide alignments (right columns) of the EL regions and the C-terminal domains of representative *ompA* variant alleles. The amino acid and nucleotide positions encompassing the EL1 to EL4 regions indicated above the sequences are those corresponding to ACICU (V1) OmpA. In VR5 the numbers at the end indicate the position of the last amino acid or nucleotide in the corresponding columns, and the high-GC 5’-GCGGCAGCTCCTGCAGCA-3’ insertion coding for the extra AAAPAA stretch found at the C-terminus in some variant sequences is shadowed in light gray. Uppercase letters in nucleotide alignments indicate the same base at a given position in all sequences. RDP4 (ref. 42) detected evidence for recombination between V3 and V4 variants at EL1 and between V4 and V5 variants at a gene region encompassing the entire EL3 and part of EL4 (shadowed in light gray). In addition, visual inspection detected sequence identity between V4 and V5 sequences along the entire EL2 (shadowed in light gray), as well as between V2 and V4 along the complete EL4 region (shadowed in dark gray only for V2). The complete amino acid and nucleotide alignments, as well as the topology predictions, are shown in Figs. S1 and S2.

### Selection acting on ompA alleles

Given that V1/V1_1 contains the largest number of *ompA* alleles, and that they are found both in *A. baumannii* and non-*baumannii* strains (Table S2), we decided to examine in detail the nature of the mutations occurring both within *A. baumannii* V1 and V1_1 alleles (19 V1 plus 2 V1_1) and also between *Acinetobacter baumannii* and the two non-*baumannii* V1_1 alleles (one from *A. seifertii* GG2 and the other from *A. nosocomialis* OIFC021) (Table S2). Within *A. baumannii* V1/V1_1 alleles (intra-species changes) the total number of synonymous substitutions was around 4-fold higher than that of non-synonymous ones (78% versus 22%) (Table 2, Figure S3), with 6/7 of the non-synonymous substitutions (86%) concentrated in the EL regions. Providing that the ELs encompass around 20% of the total length of the protein, there is a clear bias towards the EL regions carrying non-synonymous substitutions. A detailed analysis of the amino acid changes resulting from these non-synonymous substitutions indicated only conservative amino acid changes (Figure S3). Concerning inter-species changes (*A. baumannii* V1a1 *versus A. seifertii* GG2 V1_1 or *A. nosocomialis* OIFC021 V1_1, Table 2), the total number of nucleotide substitutions as compared to intra-*A. baumannii* changes increased a little depending on the non-*baumannii* species considered (40 for *A. seifertii* GG2 V1_1 and 32 for *A. nosocomialis* OIFC021 V1_1, as compared to 31 intra-species). The number of non-synonymous changes inter-species in the EL regions was similar to that found intra-species, except that one amino acid change at EL4 (Phe177 for Ala177, Figure S3) could have been positively selected. These comparisons also disclosed a substantial increase inter-species in non-synonymous substitutions in the non-exposed OmpA regions (7 in both cases as compared to none intra-species, Table 2). Moreover, 6 of these changes occurred in the TM regions, and two among them (Phe119 for Ile119 at TM5, and Tyr159 for Trp159 at TM6) are considered disfavored (Figure S3) and could have been positively selected.

**Table 2:** Summary of synonymous and non-synonymous substitutions at polymorphic sites detected between V1/V1_1 *ompA* alleles in the *Acinetobacter* population analyzed. The first 1,041 nucleotide positions (equivalent to the first 347 codons) of the different V1/V1_1 *ompA* alleles were aligned, and the mutations resulting in synonymous (syn) and non-synonymous (non-syn) changes were calculated. The numbers between brackets denote the corresponding percentages among the 347 codons in each case.

| Gene regions | Total changes | Synonymous changes | % syn | Non-synonymous changes | % non-syn |
| --- | --- | --- | --- | --- | --- |
| *a) Intra-species changes (19 V1 plus 2 V1_1 A. baumannii ompA alleles)* |  |  |  |  |  |
| All | 31 (8.9) | 24 (6.9) | 78 | 7 (2.0) | 22 |
| EL regions | 16 (4.6) | 10 (2.9) | 33 | 6 (1.7) | 19 |
| Non-exposed regions (total) | 15 (4.3) | 14 (4.0) | 45 | 1 (0.3) | 3 |
| TM regions only | 8 (2.3) | 8 (2.3) | 26 | 0 | 0 |
| *b) Inter-species changes* |  |  |  |  |  |
| *b1) A. seifertii GG2 V1_1 versus A. baumannii V1a1 ompA* |  |  |  |  |  |
| All | 40 (11.6) | 27 (7.9) | 68 | 13 (3.7) | 32 |
| EL regions | 16 (4.6) | 12 (3.4) | 29 | 4 (1.2) | 10 |
| Non-exposed regions | 24 (6.9) | 15 (4.3) | 37 | 9 (2.6) | 22 |
| TM regions only | 16 (4.6) | 9 (2.6) | 22 | 7 (2.0) | 17 |
| *b2) A. nosocomialis* OIFC021 V1_1 versus *A. baumannii* V1a1 *ompA* |  |  |  |  |  |
| All | 32 (9.2) | 20 (5.8) | 63 | 12 (3.4) | 37 |
| EL regions | 16 (4.6) | 11 (3.2) | 35 | 5 (1.4) | 15 |
| Non-exposed regions | 16 (4.6) | 9 (2.6) | 28 | 7 (2.0) | 22 |
| TM regions only | 15 (4.3) | 8 (2.3) | 25 | 7 (2.0) | 22 |

With the majority of total sequence variation within the different *A. baumannii ompA* variants located within the ELs, we hypothesized that selection may be acting differently upon these regions as compared to the rest of the protein. To test this we investigated the variation in substitution patterns for each of the V groups except for V3 and V5, as these only contained one allele each in *A. baumannii* (Table S2). For V1, V2 and V4 groups when the complete allele alignments are used, a substitution model where parameters are allowed to vary between the exposed EL regions and the non-exposed rest of the protein sequence fits the data significantly better than a fixed model where it is assumed substitution patterns are uniform across the sequence (Table S5). However, when the analysis is repeated using an alignment with recombinant regions removed, only the V2 group retains evidence for differences in substitution patterns. This suggests that, while within each V group the EL and non-exposed regions have different substitution patterns indicative of a different evolutionary history, this variation has largely been introduced by recombination.

### intra-genic recombination at the ompA locus

Detailed comparative analyses of the amino acid and corresponding nucleotide sequences spanning the *A. baumannii* OmpA variant’s variable regions (Fig. 2B) suggested a history of intra-genic recombination. The use of RDP4 (42) found evidence for intra-genic recombinatorial exchanges between V3 and V4 at EL1, and also at a longer region between V4 and V5 encompassing the entire EL3 and part of EL4 (Fig. 2B and Fig. S2). Visual inspection additionally detected substantial sequence identity between V4 and V5 along the entire EL2 and between V2 and V4 along EL4 (Fig. 2B). These observations are suggestive of several intra-genic recombinatorial exchanges during the early stages of evolution of the different *ompA* variants.

Intra-genic recombination may also explain the existence of two alternate C-terminal ends among otherwise very similar OmpA variants (see above). The main difference between them resides in the presence/absence of a hydrophobic tract of 6 amino acids composed mostly by alanine residues close to the C-terminal end (Figure 2B). We noted that the DNA sequence coding this tract, 5’-GCGGCAGCTCCTGCAGCA-3’, shows a higher GC content (72 %) when compared to the rest of the *A. baumannii ompA* gene (42 %) or even to the average GC content of *Acinetobacter* genomes (1–6). This suggests that the long QEAAAPAAAQ C-terminal region may have derived from a non-homologous (illegitimate) recombination event involving the insertion of a fragment originating in a high-GC DNA source outside the *Acinetobacter* genus (43–46). Moreover, this event is likely to have been selected recently in *A. baumannii* or in a related ACB complex species, as judged by the presence of OmpAs bearing QEAAAPAAAQ as C-terminal ends only among *A. baumannii* and other ACB species (Table S2) but not the other ecologically-differentiated clades (Table S4) into which the *Acinetobacter* genus was recently divided (1).

We analyzed possible donor(s) of the 18 nucleotide high-GC fragment among GenBank bacterial genomes by conducting a BLASTN search adjusted for short sequences identification (https://blast.ncbi.nlm.nih.gov). Among *Acinetobacter* sequences this stretch was found only in genomes of *A. baumannii* and some other ACB species, always inserted at the same position near the 3’-end of the *ompA* gene (Fig. 2A, Table S4). Remarkably however, identical or highly similar sequences(17 out of 18 nucleotide identity), which were not linked to *ompA/oprF*-related genes, were also found in the genomes of several bacterial species displaying GC contents ≥60% and assigned to different phyla including the *Actinobacteria*, the *Bacteroidetes*, and different subdivisions of the *Proteobacteria* (Table S6). Of note, the above taxa provide not only for the predominant microorganisms in soils but also for the bacterial community associated with hand palms in humans (47). These interconnecting niches, also shared by *Moraxellaceae* members including *A. baumannii* and other ACB species (1, 47), comprise a gene exchange community from which the above high-GC fragment may have been acquired by *A. baumannii* (or other ACB member) *ompA* by illegitimate recombination (44–46). The presence of identical high-GC stretches at similar gene positions in different *A. baumannii ompA* variants and subvariants (Fig. 2A) not only supports the notion that, once acquired by *ompA*, the further dissemination of this particular fragment proceeded by lateral gene transfer and fine-scale homologous recombination (43), but also that it confers some fitness advantage(s) in the human-associated niches in which *A. baumannii* and other ACB species thrive (1).

### Intra- and inter-species exchange of ompA variants

We investigated the possibility of *ompA* exchange within different lineages of the *A. baumannii* population (intra-species recombination) and also between different ACB species (inter-species recombination). The overall observations for a number of representative strains are summarized in Figure 3 (see Table S1 and Table S2 for further details). In agreement with other studies using different sets of core genes and phylogenetic strategies (1, 4, 8, 41), the *A. baumannii* strain cluster is clearly demarcated in the ML tree of Fig. 3 from those composed of its phylogenetically closely-related ACB species including *A. nosocomialis, A. seifertii, A calcoaceticus, A. oleivorans*, and *A. pittii*, which are in turn well-differentiated between them. Moreover, the *A. baumannii* strains comprising the major global clonal complexes CC1, CC2, and CC3 formed well-differentiated subclusters in this tree, as did other CCs or STs of lower representation.

**Figure 3:**
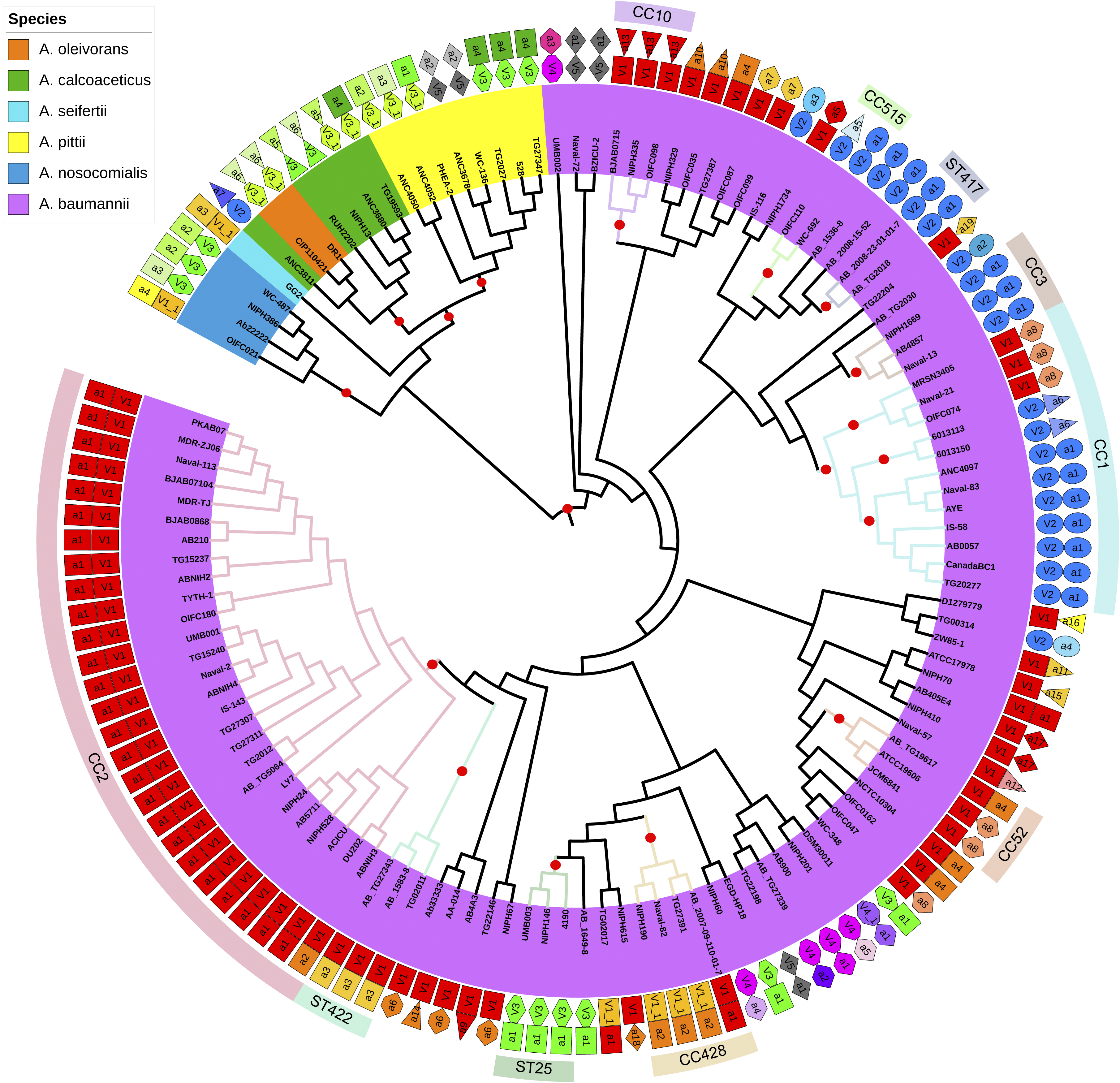
Evolutionary relatedness and *ompA* alleles of the *Acinetobacter* strains analyzed in this work. An approximately-maximum-likelihood core gene relationships was constructed using 901 core genes (present in ≥99% of strains) with the strain labels colored by species, as shown in the key. The tree is shown without scaling so that the relationships between strains can be seen, and is rooted on the branch separating the *A. baumannii* strains from the other *Acinetobacter* ACB species. Bootstrap support values ≥99% on relevant branches are indicated by red dots. The *ompA* gene alleles are indicated by the two rings of shapes surrounding the tree. The inner ring of shapes indicate the major V-group that the *ompA* allele belongs to (V1, V1_1, V2, V3, V3_1 V4, V4_1 or V5), and the outer ring indicates the specific gene allele. V1 alleles are coloured red, orange and yellow, V2 alleles are blue, V3 alleles are green, V4 alleles are pink and purple, and V5 alleles are grey. The outermost circle indicates the clusters of *A. baumannii* strains assigned to the main clonal complexes CC1, CC2, and CC3, as well as other less common clonal complexes and sequence types according to the Pasteur MLST scheme (see Table S1 for details). These CCs evolved after the middle of last century from the clonal expansion each of a highly adaptive independent genotype, and they are generally thought to represent adaptations to specific microniches and be robust with respect to gene choice (5–9).

Several additional inferences were derived from the analysis shown in Fig. 3. First, and in sharp contrast to non-*baumannii* ACB species, *ompA* V1 alleles predate the different branches of the *A. baumannii* cluster except for a few epidemiologically-related lineages (e. g., CC3, ST25, CC515 and ST417). This, added to the elevated frequency of V1 alleles among *A. baumannii* clinical strains (Table 1), and their presence already in the ATCC clinical strains 17978 and 19606 disclosed above, supports the notion that OmpA V1 variants represent long important determinants in the interaction of *A. baumannii* with the human host. Second, the distribution of *ompA* variants between different *A. baumannii* subclusters revealed non-random associations of particular variants (or variant alleles) and certain CCs (Fig. 3, see also Table S1). For instance, almost all CC2 strains (100 from 101) displayed *ompA* V1a1 with the only exception, ABNIH3, bearing V1a2 differing from V1a1 by a single synonymous substitution (Table S2). This indicated a strong pressure for the preservation of the OmpA protein codified by V1a1 during the global dissemination/expansion of CC2. In addition, V1a1 was found in two other *A. baumannii* strains branching separately from the CC2 cluster: AB_2007-09-110-01-7 and NIPH70 (Fig. 3) thus suggesting their acquisition by lateral gene transfer. AB_2007-09-110-01-7, in particular, is epidemiologically related to three other strains bearing V1_1a2 alleles (CC428, Fig. 3). This additionally supported the notion that this strain acquired V1a1 via a recombinatorial *ompA* exchange in which the donor was probably a CC2 strain. Third, the majority of the CC1 strains analyzed (26 out of 31) displayed the V2a1 allele (Fig. 3 and Table S1). These V2a1-containing strains belong to the main (L1) sublineage of CC1, estimated to have separated from the CC1 founding genotype early during the 1970s (9). This also indicates that this V2 allele was strongly preserved during the global dissemination of this CC1 sub-lineage. Two other CC1 strains (clustering together in Fig. 3) showed the V2a6 allele, differing from V2a1 by one nucleotide change translated to a conservative amino acid substitution at EL2 (Table S2). It follows that V2, and in particular V2a1, represents the predominant *ompA* variant associated with CC1. Still, the remaining three CC1 strains (which clustered together in the ML tree of Fig. 3) all showed an allele of a different variant: V1a8. Moreover, these V1a8-containing strains belong to the L2 sub-lineage of CC1 (9) (Table S1), supporting the idea of a complete *ompA* variant replacement most likely occurring very early during CC1 evolution. Fourth, similar links could be observed for other *A. baumannii* clonal lineages, as is the case of V2a1 associated with CC3, this V2 allele also with ST417 and ST417-related strains; V1a13 with CC10; V1_1a2 with ST428 and ST428-related strains; and V3a1 with ST25 and ST25-related strains (Fig. 3 and Table S1). In the case of V3a1, its presence also in strains not affiliated to ST25 such as WC-348 and EGD-HP18 (Fig. 3) provided further evidence for the existence of assortative exchanges of *ompA* variants between separate *A. baumannii* lineages.

Concerning inter-species *ompA* reassortments, evidence for exchanges between *A. baumannii* and non-*baumannii* ACB species could also be inferred from the above analysis (Fig. 3 and Table S2). For instance, V1_1 alleles were also detected in *A. nosocomialis* and *A. seifertii* strains, a V2a7 allele was found in a strain of *A. calcoaceticus*, V3 alleles were detected in *A. nosocomialis, A. calcoaceticus* and *A. pittii* strains, and a V5a2 allele was found in *A. pittii*. On the contrary, V1, V4 and V4_1 alleles were found among *A. baumannii* but not among non-*baumannii* strains and, conversely, V3_1 alleles were found in *A. oleivorans, A. calcoaceticus*, and *A. pittii* strains but not in *A. baumannii*. These differences may be related to differential role(s) of these OmpA variants in the occupation of the different niches in which the different ACB members thrive (1).

We also noted that the length of the pre-OmpA signal peptides vary between different ACB members (Table S4). Thus, pre-OmpAs of *A. baumannii, A. nosocomialis*, and *A. seifertii* all show identical signal peptides of 22 amino acid residues, while those of *A. calcoaceticus, A. oleivorans*, and *A. pittii* contain 21 amino acids. Although this could hinder in principle successful exchanges of *ompA* variants between ACB species in which the length of the signal peptide of the pre-protein differs, this seems not to be a critical limitation as judged by the presence of V2 and V3 alleles in *A. baumannii* and also in *A. calcoaceticus*, and V5 alleles in *A. baumannii* and also in *A. pittii* (Fig. 3, Table S2). These exchanges certainly required the preservation of the corresponding *ompA* signal peptide coding region in the recipient organism, and provide a further example of fine-scale intragenic recombination operating at the level of the *Acinetobacter ompA* locus, in this case following lateral gene transfer between different ACB species.

## Discussion

The study of *A. baumannii ompA* microevolution conducted here provides clues on the extent to which selection in the clinical habitat has shaped evolution of this OmpA/OprF family member. We detected five well-defined polymorphic OmpA variants in the *Acinetobacter* population analyzed, with most of the variation occurring within the exposed EL regions at the N-terminal β-barrel domain of these proteins, as well as in the C-terminal region of these proteins (Fig. 2). The EL sectors of surface-exposed bacterial proteins are subjected to strong selective pressures by many environmental factors including host defenses, and thus represent the less conserved regions with both mutation and recombination providing for the observed variabilities (23, 26, 27). Our analyses indicated that lateral gene transfer and recombination, both assortative and fine-scale, appeared as the main agents driving evolution of *A. baumannii* OmpA. We found signs of different fine-scale intra-genic recombination events in *A. baumannii ompA* variants, some encompassing the EL regions which appeared to have occurred during their early stages of divergence from their recent common ancestor. Other events, which appear more subtle and more recent, resulted in different C-terminal sequences in the otherwise highly conserved C-terminal periplasmic domain of OmpA (Fig. 2). The existence of two alternate C-terminal ends differing in an alanine-rich AAAPAA tract not only among different OmpA variants but also within a given variant, as in the case of V1 and V4 (Fig. 2), strongly point to lateral gene transfer and fine-scale homologous recombination affecting the conserved *ompA* 3’-region as the likely cause of this variability. Our analysis additionally suggested that the high-GC DNA fragment encoding this tract represents a recent acquisition resulting from illegitimate recombination occurring in an *A. baumannii* (or a closely-related ACB species) member adapting to human-associated environments. *Acinetobacter* members can take up DNA from environmental sources including soil microcosms, and incorporate short fragments into the genome by homologous as well as non-homologous recombination (44–46). In fact, natural transformation with highly fragmented (≥ 20 bp) and even partially damaged DNA has been shown to be much more efficient than spontaneous mutation to generate adjacent nucleotide polymorphism in the *Acinetobacter* genome (46). The OmpA periplasmic domain contributes to the stability of the cell envelope by interacting non-covalently with the peptidoglycan (25), and its deletion in *A. baumannii* resulted in an increased cell susceptibility to different antimicrobials including cell wall synthesis inhibitors (50). This region of OmpA has been found to associate with different OM and periplasmic proteins including an acquired OXA-23 β-lactamase in the carbapenem-resistant *A. baumannii* strain AB5075 (36). It is then tempting to speculate that the insertion of the hydrophobic Ala tract at the OmpA C-terminal end may have resulted in an increased envelope stability towards the various damaging substances used in the nosocomial environment (29, 48, 49), thus providing the driving force for its selection during adaptation of *A. baumannii* to this particular niche.

Analyses of the substitution patterns in the V1 and V1_1 *ompA* alleles provided further evidence for selection acting upon the protein (Table 2). When looking within *A. baumannii*, non-synonymous substitutions were concentrated in the EL regions of the sequence, suggesting adaptation to the extracellular environment. However, when comparing between *A. baumannii* and other ACB species, more non-synonymous changes were found in the TM regions. If we consider that non-synonymous substitutions provide a measure of adaptive changes to a given environment, then recombinatorial exchanges of V1/V1_1 *ompA* alleles between *A. baumannii* and non-*baumannii* species were apparently followed by the rapid selection of adaptive mutations mostly at the level of the TM sectors in the receiving species. This may have resulted from different physicochemical properties of the OM between *A. baumannii* and non-*baumannii* species, and offers another example that the boundaries between ACB species may limit, but not obliterate, inter-species lateral transfer of *ompA* alleles.

The different OmpA variants were unequally distributed among the *A. baumannii* population, with V1 alleles being found in around two thirds of the strains analyzed and V2 alleles in around one fourth of them (Table 1). We found a strong pressure for the preservation of particular alleles of these two variants during the dissemination and expansion of the major *A. baumannii* global clones CC1 and CC2. Thus, V1a1 *ompA* was found in almost all CC2 strains, while V2a1 was predominant among the main L1 lineage of CC1 (Fig. 3). This suggests that the characteristics evolved by these two OmpA variants confer significant fitness advantages to the strains that compose these epidemic *A. baumannii* clonal complexes. Significant genetic diversity is seemingly a hallmark of *A. baumannii*, both between epidemiologically- and phylogenetically-closely related strains (2, 3, 5, 9, 11, 12). In both CC2 and CC1 strains different whole-genome sequencing studies have indicated that most of this diversity is driven by recombination, with genetic material derived from different co-existing *A. baumannii* lineages (5, 9, 11, 12). Many of these exchanges induced loss or swapping-out of exposed structures potentially recognized by host defenses, such as the capsular polysaccharide, the outer core lipooligosaccharide, iron capture components, efflux pumps, and OM proteins such as CarO (9–12). It is generally assumed that this ongoing process contributes to immune evasion facilitating long-term patient colonization and spread of the *A. baumannii* epidemic clones (9–12). *A. baumannii* OmpA has been found capable of eliciting immune responses including the production of anti-OmpA antibodies both in infected patients (17, 37) and experimentally infected mice (15, 20). It is then noteworthy that, in sharp contrast with the other surface-exposed macromolecules mentioned above, particular *ompA* V1 and V2 alleles have been strongly preserved during the dissemination and expansion of the main *A. baumannii* global clones CC1 and CC2. This preservation supports the notion that these two OmpA variants play pivotal roles in *A. baumannii* pathogenicity (3, 6, 18, 19, 24, 28–36). Their escape from host defenses could result, at least in part, from the recently disclosed shielding by capsule polysaccharide production during the course of *A. baumannii* infection (50). Thus, the observation that particular *ompA* variants, variant alleles, or even *ompA* gene sectors can be effectively exchanged by either assortative or fine-scale recombination not only among different *A. baumannii* lineages but also between different ACB species (Fig. 3) represents a worrying scenario, providing that any innovation evolved by *ompA* affecting host specificity or persistence may be rapidly disseminated among all components of this expanded *Acinetobacter* community.

Finally, there has been interest in developing a vaccine against *A. baumannii* using OmpA as immunogen (14–20, 37). However, the recombinatorial potential of *ompA* variants shown in this work may pose severe limitations to the effectiveness of a vaccine based solely on a single OmpA variant. It remains to be tested whether polyvalent OmpA vaccines generated through directed protein evolution (51) using the analysis provided in this work could prevent infections due to *A. baumannii* and other ACB strains bearing different *ompA* variants. Further investigations involving functional studies of different OmpA variants/variant alleles may shed light upon the relative strengths of selection acting upon these OM proteins and help the development of new strategies for combating *A. baumannii* and other ACB infections.

## Materials and Methods

### Sequence retrieval, alignment and phylogeny estimation

The nucleotide sequence of *ompA* from *A. baumannii* ACICU (accession number NC_017162.1, locus tag ABK1_RS15935) was used to query the JGI SIMG/M database (https://img.jgi.doe.gov/) for available *ompA* sequences from *Acinetobacter* genomes. Nucleotide sequences were translated, aligned, and separated into groups of similarity. Based on these groupings, an alignment of translated *ompA* sequences of the 21 identified *A. baumannii* OmpA protein alleles was used to estimate an unrooted maximum-likelihood (ML) phylogenetic tree using PhyML (http://phylogeny.lirmm.fr/phylo_cgi/index.cgi). The assembled *Acinetobacter* strain genomes used were obtained from the NCBI database, and their MLST types (Pasteur scheme) were retrieved from https://pubmlst.org/bigsdb?db=pubmlst_abaumannii_isolates&page=query. A core gene alignment was obtained using Roary (52), and the ML phylogeny was estimated using FastTree v2.1.10, with support estimated from 1,000 resamples (53). For *Acinetobacter* strains where the species designation was uncertain, *rpoB* sequences were used in a BLAST search to confirm the species to which they belong (54).

### Protein sequence diversity

Variation between *A. baumannii* OmpA variants was determined by calculating the Shannon entropy from alignments of the 21 alleles described here (https://www.hiv.lanl.gov/content/sequence/ENTROPY/entropy.html), as described previously for *A. baumannii* CarO (10).

### Intragenic recombination detection

An alignment containing one of each *A. baumannii ompA* allele was loaded into the Recombination Detection Program v4 (RDP4) (42). An automated recombination analysis was run using the RDP (55), GENCONV (56) and Chimera (57) methods. Regions in which significant evidence for recombination was detected were removed using the ‘Save alignment with recombinant regions removed’ feature in RDP, and used in subsequent analyses.

### Tests for selection

To investigate whether natural selection is acting similarly in the *A. baumannii ompA* EL regions as compared to the rest of the protein, we used a partition model in the program codeml implemented in PAML v4.9d (58, 59). Separate alignments for alleles from groups V1, V2 and V4 either including all data, or with recombinant regions removed, were used to estimate trees using PhyML, which then had distance information manually removed for use as input to codeml. Input alignment files were partitioned into sequences corresponding to exposed EL regions and those corresponding to non-exposed TM and periplasmic domains. Two models were fitted to the data in codeml, a null model where parameters values are assumed to be equal between the exposed and non-exposed regions of the sequence (model = 0, NSsites = 0, Mgene = 0), and an alternative model where *k* (transition/transversion rate ratio), *ω* (*d*_N_/*d*_S_), *π*s (codon frequencies) and *c*s (proportional branch lengths) are allowed to vary between the two regions (model = 0, NSsites = 0, Mgene = 4). The resulting log likelihood values from the two models were compared using χ^2^ tests to determine whether the alternative models better fit the data than the null models. V3 and V5 were excluded from this analysis as they only contain one allele each in *A. baumannii*.

