## Supplementary material for "Microevolution in the major outer membrane protein OmpA of *Acinetobacter baumannii*": Figure S1

Figure S1: Protein sequence comparisons and predicted outer membrane topology of the different *A. baumannii* OmpA variants. Alignments were conducted using ClustalW (http://www.genome.jp/tools/clustalw/) and refined by visual inspection. The *A. baumannii* isolates bearing the corresponding representative variants are indicated in the left column. The numbers at the right indicate the position of the last aa residue for the corresponding variant in a particular row. The symbols below the alignments indicate identical (*) or conserved (:) aa residues at a given position, deletions are indicated in the sequences by hyphens (-). Topology predictions were done using PRED-TMBB (http://bioinformatics.biol.uoa.gr/PRED-TMBB/), N-terminal transit peptides are highlighted in yellow, periplasmic regions in green, transmembrane (TM) regions in red, and external loops (EL) in light blue. The 20 aa stretch bearing the **N**$X_2$**LS**$X_2$**RA**$X_2$**V**$X_2$**L** conserved motif at the C-terminal domain (underlined with " α" symbols) points to the predicted α-helix interacting with the peptidoglycan common to OmpA-like domain proteins (25). Note also the 6 aa indel (indicated in magenta) near the C-terminal region resulting in two alternate C-terminal ends of different lengths and hydrophobicity.

```
                            Transit peptide           TM1            EL1
ACICU_V1_a1             MKLSRIALATMLVAAPLAAANAGVTVTPLLLGYTFQDSQHNNGGKDGNLTNSPELQDDLF  60
TG02017_V1_1_a1         MKLSRIALATMLVAAPLAAANAGVTVTPLLLGYTFQDSQHNNGGKDGNLTNAPELQDDLF  60
AB0057_V2_a_1           MKLSRIALATMLVAAPLAAANAGVTVTPLLLGYTFQDTQHNNGGKDGELTNGPELQDDLF  60
4190_V3_a_1             MKLSRIALATMLVAAPLAAANAGVTVTPLLLGYTWQDSEHNN----NKLTDHAELQDDLF  56
AB900_V4_a_1            MKLSRIALATMLVAAPLAAANAGVTVTPLLLGYTFQDSEHNN----HKLTDSPELQDDLF  56
DSM30011_V4_1_a_1       MKLSRIALATMLVAAPLAAANAGVTVTPLLLGYTFQDSEHNN----HKLTDSPELQDDLF  56
BZICU-2_V5_a_1          MKLSRIALATMLVAAPLAAANAGVTVTPLLLGYTFQDSQHNNGGKDGSLTNGPELQDDLF  60
                        **********************************:**::***      **: :*******

                          TM2          TM3          EL2              TM4          TM5
ACICU_V1_a1             VGAALGIELTPWLGFEAEYNQVKGDVDGASA-GAEYKQKQINGNFYVTSDLITKNYDSKI 119
TG02017_V1_1_a1         VGAALGIELTPWLGFEAEYNQVKGDVDGASA-GAEYKQKQINGNFYVTSDLITKNYDSKI 119
AB0057_V2_a_1           VGAALGIELTPWLGFEAEYNQVKGDVDGLAA-GAEYKQKQINGNFYVTSDLITKNYDSKI 119
4190_V3_a_1             VGAGLGVELTPWLGFEAEYNQVKGDLDGTGVQGAEYKQKTIAGNFYATSDLITKNYDSKF 116
AB900_V4_a_1            VGAALGIELTPWLGFEAEYNQVKGDVD---TNYGEYKQKQINGNFYVTSDLITKNYDSKI 113
DSM30011_V4_1_a_1       VGAALGIELTPWLGFEAEYNQVKGDVD---PNYGEYKQKQINGNFYVTSDLITKNYDSKI 113
BZICU-2_V5_a_1          VGAALGIELTPWLGFEAEYNQVKGDVD---TNYGEYKQKQINGNFYVTSDLITKNYDSKI 117
                        *** **:******************:*       ***** * **** ************:

                        TM5              EL3                 TM6            TM7  EL4
ACICU_V1_a1             KPYVLLGAGHYKYDFDGVNRGTRGTSEEGTLGNAGVGAFWRLNDALSLRTEARATYNADE 179
TG02017_V1_1_a1         KPYVLLGAGHYKYDFDGVNRGTRGTSEEGTLGNAGVGAFWRLNDALSLRTEARATYNADE 179
AB0057_V2_a1            KPYVLLGAGHYKYEIPDL---SYHNDEEGTLGNAGVGAFWRLNDALSLRTEARGTYNFDE 176
4190_V3_a1              KPYVLLGAGQTKTEFDGI-----YEDKKDTIGNAGVGAFYRLNDALSLRTEARGTYDFDE 171
AB900_V4_a1             KPYVLLGAGHYKYDFDDA-RLAYHDGEEGTLGNAGVGAFWRLNDALSLRTEARGTYNFDE 172
DSM30011_V4_1_a1        KPYVLLGAGHYKYDFDDA-RLAYHDGEEGTLGNAGIGAFWRLNDALSLRTEARGTYNFDE 172
BZICU-2_V5_a1           KPYVLLGAGHYKYDFDDA-RLAYHDGEEGTLGNAGVGAFWRLNDALSLRTEARGTYNFDE 176
                        *********: * ::           :: *:****:***:************* **: **

                        EL4     TM8
ACICU_V1_a1             EFWNYTALAGLNVVLGGHLKPAAPVVEVAPVEPTPVAPQPQELTEDLNMELRVFFDTNKS 239
TG02017_V1_1_a1         EFWNYTALAGLNVVLGGHLKPAAPVVEVAPVEPTPVAPQPQELTEDLNMELRVFFDTNKS 239
AB0057_V2_a_1           KFWNYTALAGLNVVLGGHLKPAAPVVEVAPVEPTPVAPQPQELTEDLNMELRVFFDTNKS 236
4190_V3_a_1             KYWRYTALAGLNVVLGGHLKPAAPVVEVAPVEPTPVAPQPQELTEDLNMELRVFFDTNKS 231
AB900_V4_a_1            KFWNYTALAGLNVVLGGHLKPAAPVVEVAPVEPTPVAPQPQELTEDLNMELRVFFDTNKS 232
DSM30011_V4_1_a_1       KFWNYTALAGLNVVLGGHLKPAAPVVEVAPVEPTPVAPQPQELTEDLNMELRVFFDTNKS 232
BZICU-2_V5_a_1          QFWNYTALAGLNVVLGGHLKPAAPVVEVAPVEPTPVAPQPQELTEDLNMELRVFFDTNKS 236
                        ::* ********************************************************

ACICU_V1_a1             NIKDQYKPEIAKVAEKLSEYPNATARIEGHTDNTGPRKLNERLSLARANSVKSALVNEYN 299
TG02017_V1_1_a1         NIKDQYKPEIAKVAEKLSEYPNATARIEGHTDNTGPRKLNERLSLARANSVKSALVNEYN 299
AB0057_V2_a_1           NIKDQYKPEIAKVAEKLSEYPNATARIEGHTDNTGPRKLNERLSLARANSVKSALVNEYN 296
4190_V3_a_1             NIKDQYKPEIAKVAEKLSEYPNATARIEGHTDNTGPRKLNERLSLARANSVKSALVNEYN 291
AB900_V4_a_1            NIKDQYKPEIAKVAEKLSEYPNATARIEGHTDNTGPRKLNERLSLARANSVKSALVNEYN 292
DSM30011_V4_1_a_1       NIKDQYKPEIAKVAEKLSEYPNATARIEGHTDNTGPRKLNERLSLARANSVKSALVNEYN 292
BZICU-2_V5_a_1          NIKDQYKPEIAKVAEKLSEYPNATARIEGHTDNTGPRKLNERLSLARANSVKSALVNEYN 296
                        ************************************************************

                                                            ααααααααααααααααααααααα

ACICU_V1_a1             VDASRLSTQGFAWDQPIADNKTKEGRAMNRRVFATITGSRTVVVQPGQEAAAPAAAQ 356
TG02017_V1_1_a1         VDASRLSTQGFAWDQPIADNKTKEGRAMNRRVFATITGSRTVVVQPGQQ------AQ 350
AB0057_V2_a_1           VDASRLSTQGFAWDQPIADNKTKEGRAMNRRVFATITGSRTVVVQPGQEAAAPAAAQ 353
4190_V3_a_1             VDASRLSTQGFAWDQPIADNKTKEGRAMNRRVFATITGSRTVVVQPGQQ------AQ 342
AB900_V4_a_1            VDASRLSTQGFAWDQPIADNKTKEGRAMNRRVFATITGSRTVVVQPGQEAAAPAAAQ 349
DSM30011_V4_1_a_1       VDASRLSTQGFAWDQPIADNKTKEGRAMNRRVFATITGSRTVVVQPGQQ------AQ 343
BZICU-2_V5_a_1          VDASRLSTQGFAWDQPIADNKTKEGRAMNRRVFATITGSRTVVVQPGQQ------AQ 347
                        ************************************************:      **
```
