## Supplementary material for "Microevolution in the major outer membrane protein OmpA of *Acinetobacter baumannii*": Figure S2

Figure S2: Nucleotide sequence comparisons of *A. baumannii ompA* variants. DNA alignments were done using ClustalW (http://www.genome.jp/tools/clustalw/) and refined by visual inspection on the basis of the protein alignments of Fig. S1. As in the previous Figure, the *A. baumannii* isolates bearing representative *ompA* variant genes are indicated at the left (DSM denotes DSM30011), and the numbers at the right indicate the position of the last nt residue of the corresponding *ompA* variant gene in a particular row. Uppercase letters denote the same nt in all sequences at a given position. Deletions are indicated by hyphens (-). The 18-nt indel and the C per G substitution near the TAA end codon resulting in variable OmpA C-terminal motifs are indicated in magenta. The nt sections coding for the N-terminal transit peptides, periplasmic regions, transmembrane (TM) regions, and external loops (EL) are highlighted as in Fig. S1 above. Evidence for recombination  was found by using the RDP4 sofware (42) between AB900/DSM30011 (V4, major parent) and 4190 (V3, minor parent) at the EL1 region encompassing nt positions 121 to 141 of these sequences (4 programs out of 7, shadowed in light gray). RDP4 also indicated recombination (6 programs out of 7) between AB900/DSM30011 (V4) and BZICU-2 (V5) at a gene region spanning nt positions 343 (V4)/355 (V5) to 512 (V4)/524 (V5), which encompasses the entire EL3 and part of EL4 (shadowed in gray).

```
                                          Transit peptide coding region
ACICU_V1_a1       ATGAAATTGAGTCGTATTGCACTTGCTACTATGCTTGTTGCTGCTCCATTAGCTGCTGCT   60
TG02017_V1_1_a1   ATGAAATTGAGTCGTATTGCACTTGCTACTATGCTTGTTGCTGCTCCATTAGCTGCTGCT   60
AB0057_V2_a1      ATGAAATTGAGTCGTATTGCACTTGCTACTATGCTTGTTGCTGCTCCATTAGCTGCTGCT   60
4190_V3_a1        ATGAAATTGAGTCGTATTGCACTTGCTACTATGCTTGTTGCTGCTCCATTAGCTGCTGCT   60
AB900_V4_a1       ATGAAATTGAGTCGTATTGCACTTGCTACTATGCTTGTTGCTGCTCCATTAGCTGCTGCT   60
DSM_V4_1_a1       ATGAAATTGAGTCGTATTGCACTTGCTACTATGCTTGTTGCTGCTCCATTAGCTGCTGCT   60
BZICU-2_V5_a1     ATGAAATTGAGTCGTATTGCACTTGCTACTATGCTTGTTGCTGCTCCATTAGCTGCTGCT   60

                                                   TM1                    EL1
ACICU_V1_a1       AATGCTGGCGTAACAGTTACTCCATTAcTGCTTGGTTAcACtTtcCAAGAcagccAaCAc  120
TG02017_V1_1_a1   AATGCTGGCGTAACAGTTACTCCATTATTGCTTGGTTAcACtTtcCAAGAcagccAaCAc  120
AB0057_V2_a1      AATGCTGGCGTAACAGTTACTCCATTATTGCTTGGTTAcACtTtcCAAGAcactcAaCAc  120
4190_V3_a1        AATGCTGGCGTAACAGTTACTCCATTATTGCTTGGTTAtACtTggCAAGAcagcgAgCAc  120
AB900_V4_a1       AATGCTGGCGTAACAGTTACTCCATTATTGCTTGGTTAcACaTtcCAAGAttctgAaCAt  120
DSM_V4_1_a1       AATGCTGGCGTAACAGTTACTCCATTATTGCTTGGTTAcACaTtcCAAGAttctgAaCAt  120
BZICU-2_V5_a1     AATGCTGGCGTAACAGTTACTCCATTATTGCTTGGTTAcACtTtcCAAGAcagccAaCAc  120

                                     EL1
ACICU_V1_a1       AACAAtggcggtaaagatggtaacTTaACtaAttcacCtGAgtTaCAAGAcGATTTATTC  180
TG02017_V1_1_a1   AACAAtggcggtaaagatggtaacTTaACtaAtgcacCtGAgtTaCAAGAcGATTTATTC  180
AB0057_V2_a1      AACAAtggcggtaaagatggcgagTTaACtaAcggacCtGAatTaCAAGAcGATTTATTC  180
4190_V3_a1        AACAAc------------aataaaTTaACtgAtcatgCtGAatTgCAAGAtGATTTATTt  168
AB900_V4_a1       AACAAc------------cataaaTTaACtgAtagccCaGAgcTaCAAGAtGATTTATTt  168
DSM_V4_1_a1       AACAAc------------cataaaTTaACtgAtagccCaGAgcTaCAAGAtGATTTATTt  168
BZICU-2_V5_a1     AACAAtggcggtaaagatggtagcTTgACaaAtggtcCtGAgtTaCAAGAcGATTTATTC  180

                      TM2                                TM3
ACICU_V1_a1       GTTGGtGCaGcTCTgGGTaTcGAGTTAACtCCTTGGTTAGGTTTcGAAGCTGAATATAAc  240
TG02017_V1_1_a1   GTTGGcGCaGcTCTgGGTaTcGAGTTAACaCCTTGGTTAGGTTTcGAAGCTGAATATAAc  240
AB0057_V2_a1      GTTGGtGCaGcTCTTGGTaTcGAGTTAACaCCTTGGTTAGGTTTcGAAGCTGAATATAAc  240
4190_V3_a1        GTTGGtGCtGgTCTgGGTgTtGAGTTAACaCCTTGGTTAGGTTTtGAAGCTGAATATAAt  228
AB900_V4_a1       GTTGGtGCaGcTCTtGGTaTcGAGTTAACaCCTTGGTTAGGTTTcGAAGCTGAATATAAc  228
DSM_V4_1_a1       GTTGGtGCaGcTCTtGGTaTcGAGTTAACaCCTTGGTTAGGTTTcGAAGCTGAATATAAc  228
BZICU-2_V5_a1     GTTGGtGCaGcTCTtGGTaTcGAGTTAACaCCTTGGTTAGGTTTtGAAGCTGAATATAAc  240

                                     EL2
ACICU_V1_a1       CAAGTtAAAGGcGAcgTaGAcggcgcttctgct---ggtGcTGAaTAtAAaCAaAAacaa  297
TG02017_V1_1_a1   CAAGTtAAAGGtGAcgTaGAcggcgcttctgct---ggtGcTGAaTAtAAaCAaAAacaa  297
AB0057_V2_a1      CAAGTtAAAGGtGAtgTaGAcggtcttgcagct---ggcGcTGAaTAcAAgCAaAAacaa  297
4190_V3_a1        CAAGTaAAAGGtGAtcTtGAtggtactggcgttcaaggcGcTGAgTAcAAaCAgAAgact  288
AB900_V4_a1       CAAGTtAAAGGtGAtgTaGAc---------acgaactatGgTGAaTAtAAaCAgAAacaa  279
DSM_V4_1_a1       CAAGTtAAAGGtGAtgTaGAt---------ccaaactacGgTGAaTAtAAgCAgAAacaa  279
BZICU-2_V5_a1     CAAGTtAAAGGtGAtgTaGAc---------acgaactatGgTGAaTAcAAaCAaAAacaa  291

                         TM4                                        TM5
ACICU_V1_a1       ATcaacGGTAACTTCTATGttACTTCTGATTTAATtACTAAAAACTAcGAcAGCAAAaTC  357
TG02017_V1_1_a1   ATcaacGGTAACTTCTATGttACTTCTGATTTAATtACTAAAAACTAcGAcAGCAAAaTC  357
AB0057_V2_a1      ATcaacGGTAACTTCTATGttACTTCTGATTTAATcACTAAAAACTAtGAcAGCAAAaTC  357
4190_V3_a1        ATtgctGGTAACTTCTATGcaACTTCTGATTTAATcACTAAAAACTAtGAcAGCAAAtTt  348
AB900_V4_a1       ATcaacGGTAACTTCTATGttACTTCTGATTTAATtACTAAAAACTAcGAtAGCAAAaTC  339
DSM_V4_1_a1       ATcaatGGTAACTTCTATGttACTTCTGATTTAATtACTAAAAACTAcGAtAGCAAAaTC  339
BZICU-2_V5_a1     ATcaaCGGTAACTTCTATGttACTTCTGATTTAATtACTAAAAACTAtGAcAGCAAAaTC  351

                                                         EL3
ACICU_V1_a1       AAgCCgTAcGTATTaTTAGGTGCtGGTCActatAAAtatGActTTgaTGgcgtaaatcgt  417
TG02017_V1_1_a1   AAgCCgTAcGTATTaTTAGGTGCtGGTCActatAAAtacGActTTgaTGgcgtaaaccgt  417
AB0057_V2_a1      AAgCCaTAcGTATTgTTAGGTGCtGGTCActacAAAtacGAgaTTccTGacctt------  411
4190_V3_a1        AAgCCaTAcGTATTgTTAGGTGCgGGTCAaacTAAAactGAgtTTgaTGgtatc------  402
AB900_V4_a1       AAaCCtTAcGTATTgTTAGGTGCaGGTCAttatAAAtatGAttTTgaTGatgct---cgt  396
DSM_V4_1_a1       AAaCCtTAcGTATTgTTAGGTGCaGGTCAttatAAAtatGAttTTgaTGatgct---cgt  396
BZICU-2_V5_a1     AAgCCtTAtGTATTgTTAGGTGCaGGTCAttatAAAtatGAttTTgaTGatgct---cgt  408

                                                           TM6
ACICU_V1_a1       ggtacacgtggtacttctgAagAAGgTACTtTaGGTAAcGCTGGTgTtGGTGCTTTCTgg  477
TG02017_V1_1_a1   ggtaCacgtggtacttctgAagAAGgTACTtTaGGTAAcGCTGGTgTtGGTGCTTTCTgg  477
AB0057_V2_a1      ---tcttatcacaacgatgAagAAGgTACTtTaGGTAAtGCTGGTgTtGGTGCTTTCTgg  468
4190_V3_a1        ---------tatgaagacaAgaAAGaTACTaTcGGTAAtGCcGGTgTaGGTGCTTTCTat  453
```

```
AB900_V4_a1        ttagcttaccatgatggtgAagAAGgTACTtTaGGTAAcGCTGGTgTtGGTGCTTTCTgg  456
DSM_V4_1_a1        ttagcttaccatgatggtgAagAAGgTACTtTaGGTAAcGCTGGTaTtGGTGCTTTCTgg  456
BZICU-2_V5_a1      ttagcttaccatgatggtgAagAAGgTACTtTaGGTAAtGCTGGTgTtGGTGCTTTCTgg  468

                                                TM7                     EL4
ACICU_V1_a1        CGCTTaAAcGAcGCTtTaTCTCTTCGTACtGAAGCTCGtGcTACtTATaAtgcTGAtGAA  537
TG02017_V1_1_a1    CGCTTaAAcGAcGCTtTaTCTCTTCGTACtGAAGCTCGtGcTACtTATaAtgcTGAtGAA  537
AB0057_V2_a1       CGCTTaAAcGAtGCTcTaTCTCTTCGTACaGAAGCTCGtGgTACtTATaActtTGAcGAA  528
4190_V3_a1         CGCTTgAAcGAtGCTTTgTCTCTTCGTACaGAAGCTCGcGgTACgTATgAtttTGAtGAA  513
AB900_V4_a1        CGCTTaAAtGAtGCTtTaTCTCTTCGTACaGAAGCTCGtGgTACtTATaActtTGAcGAA  516
DSM_V4_1_a1        CGCTTaAAcGAtGCTtTaTCTCTTCGTACaGAAGCTCGtGgTACtTATaActtTGAcGAA  516
BZICU-2_V5_a1      CGCTTaAAtGAcGCTtTaTCTCTTCGTACaGAAGCgCGtGgTACtTATaActtTGAtGAA  528

                                               TM8
ACICU_V1_a1        gAgTtCTGGaaCTAtACaGCtCTTGCTGGCTTAAACGTAGTTCTTGGTGGTCACTTGAAG  597
TG02017_V1_1_a1    gAgTtCTGGaaCTAtACaGCtCTTGCTGGCTTAAACGTAGTTCTTGGTGGTCACTTGAAG  597
AB0057_V2_a1       aAaTtCTGGaaCTAtACaGCtCTTGCTGGCTTAAACGTAGTTCTTGGTGGTCACTTGAAG  588
4190_V3_a1         aAaTaCTGGcgCTAcACtGCtCTTGCTGGCTTAAACGTAGTTCTaGGTGGTCACTTGAAG  573
AB900_V4_a1        aAaTtCTGGaaCTAtACaGCtCTTGCTGGCTTAAACGTAGTTCTTGGTGGTCACTTGAAG  576
DSM_V4_1_a1        aAaTtCTGGaaCTAtACaGCaCTTGCTGGCTTAAACGTAGTTCTTGGTGGTCACTTGAAG  576
BZICU-2_V5_a1      cAaTtCTGGaaCTAtACaGCtCTTGCTGGCTTAAACGTAGTTCTTGGTGGTCACTTGAAG  588

ACICU_V1_a1        CCTGCTGCTCCTGTAGTAGAAGTTGCTCCAGTTGAACCAACTCCAGTTGCTCCACAACCA  657
TG02017_V1_1_a1    CCTGCTGCTCCTGTAGTAGAAGTTGCTCCAGTTGAACCAACTCCAGTTGCTCCACAACCA  657
AB0057_V2_a1       CCTGCTGCTCCTGTAGTAGAAGTTGCTCCAGTTGAACCAACTCCAGTTGCTCCACAACCA  648
4190_V3_a1         CCTGCTGCTCCTGTAGTAGAAGTTGCTCCAGTTGAACCAACTCCAGTTGCTCCACAACCA  633
AB900_V4_a1        CCTGCTGCTCCTGTAGTAGAAGTTGCTCCAGTTGAACCAACTCCAGTTGCTCCACAACCA  636
DSM_V4_1_a1        CCTGCTGCTCCTGTAGTAGAAGTTGCTCCAGTTGAACCAACTCCAGTTGCTCCACAACCA  636
BZICU-2_V5_a1      CCTGCTGCTCCTGTAGTAGAAGTTGCTCCAGTTGAACCAACTCCAGTTGCTCCACAACCA  648

ACICU_V1_a1        CAAGAGTTAACTGAAGACCTTAACATGGAACTTCGTGTgTTCTTTGATACTAACAAATCA  717
TG02017_V1_1_a1    CAAGAGTTAACTGAAGACCTTAACATGGAACTTCGTGTgTTCTTTGATACTAACAAATCA  708
AB0057_V2_a1       CAAGAGTTAACTGAAGACCTTAACATGGAACTTCGTGTgTTCTTTGATACTAACAAATCA  693
4190_V3_a1         CAAGAGTTAACTGAAGACCTTAACATGGAACTTCGTGTgTTCTTTGATACTAACAAATCA  717
AB900_V4_a1        CAAGAGTTAACTGAAGACCTTAACATGGAACTTCGTGTaTTCTTTGATACTAACAAATCA  696
DSM_V4_1_a1        CAAGAGTTAACTGAAGACCTTAACATGGAACTTCGTGTgTTCTTTGATACTAACAAATCA  708
BZICU-2_V5_a1      CAAGAGTTAACTGAAGACCTTAACATGGAACTTCGTGTgTTCTTTGATACTAACAAATCA  696

ACICU_V1_a1        AACATCAAAGACCAaTACAAGCCAGAAATcGCTAAAGTTGCTGAAAAATTATCTGAATAC  777
TG02017_V1_1_a1    AACATCAAAGACCAaTACAAGCCAGAAATcGCTAAAGTTGCTGAAAAATTATCTGAATAC  777
AB0057_V2_a1       AACATCAAAGACCAaTACAAGCCAGAAATcGCTAAAGTTGCTGAAAAATTATCTGAATAC  768
4190_V3_a1         AACATCAAAGACCAgTACAAGCCAGAAATcGCTAAAGTTGCTGAAAAATTATCTGAATAC  753
AB900_V4_a1        AACATCAAAGACCAaTACAAGCCAGAAATtGCTAAAGTTGCTGAAAAATTATCTGAATAC  756
DSM_V4_1_a1        AACATCAAAGACCAaTACAAGCCAGAAATcGCTAAAGTTGCTGAAAAATTATCTGAATAC  756
BZICU-2_V5_a1      AACATCAAAGACCAaTACAAGCCAGAAATtGCTAAAGTTGCTGAAAAATTATCTGAATAC  768

ACICU_V1_a1        CCTAACGCTACTGCACGTATCGAAGGTCAcACAGATAACACTGGTCCACGTAAGTTGAAC  837
TG02017_V1_1_a1    CCTAACGCTACTGCACGTATCGAAGGTCAcACAGATAACACTGGTCCACGTAAGTTGAAC  837
AB0057_V2_a1       CCTAACGCTACTGCACGTATCGAAGGTCAcACAGATAACACTGGTCCACGTAAGTTGAAC  828
4190_V3_a1         CCTAACGCTACTGCACGTATCGAAGGTCAcACAGATAACACTGGTCCACGTAAGTTaAAC  813
AB900_V4_a1        CCTAACGCTACTGCACGTATCGAAGGTCAcACAGATAACACTGGTCCACGTAAGTTGAAC  816
DSM_V4_1_a1        CCTAACGCTACTGCACGTATCGAAGGTCAtACAGATAACACTGGTCCACGTAAGTTGAAC  816
BZICU-2_V5_a1      CCTAACGCTACTGCACGTATCGAAGGTCAcACAGATAACACTGGTCCACGTAAGTTGAAC  828

ACICU_V1_a1        GAACGTTTATCTTTAGCTCGTGCTAACTCTGTTAAATCAGCTCTTGTAAACGAATAcAAC  897
TG02017_V1_1_a1    GAACGTTTATCTTTAGCTCGTGCTAACTCTGTTAAATCAGCTCTTGTAAACGAATAtAAC  897
AB0057_V2_a1       GAACGTTTATCTTTAGCTCGTGCTAACTCTGTTAAATCAGCTCTTGTAAACGAATAcAAC  888
4190_V3_a1         GAACGTTTATCTTTAGCTCGTGCTAACTCTGTTAAATCAGCTCTTGTAAACGAATAcAAC  873
AB900_V4_a1        GAACGTTTATCTTTAGCTCGTGCTAACTCTGTTAAATCAGCTCTTGTAAACGAATAcAAC  876
DSM_V4_1_a1        GAACGTTTATCTTTAGCTCGTGCTAACTCTGTTAAATCAGCTCTTGTAAACGAATAtAAC  876
BZICU-2_V5_a1      GAACGTTTATCTTTAGCTCGTGCTAACTCTGTTAAATCAGCTCTTGTAAACGAATAtAAC  888

ACICU_V1_a1        GTTGACGCTTCTCGTTTGTCTACTCAAGGTTTCGCTTGGGATCAACCGATTGCTGACAAC  957
TG02017_V1_1_a1    GTTGACGCTTCTCGTTTGTCTACTCAAGGTTTCGCTTGGGATCAACCGATTGCTGACAAC  957
AB0057_V2_a1       GTTGACGCTTCTCGTTTGTCTACTCAAGGTTTCGCTTGGGATCAACCGATTGCTGACAAC  948
4190_V3_a1         GTTGACGCTTCTCGTTTGTCTACTCAAGGTTTCGCTTGGGATCAACCGATTGCTGACAAC  933
AB900_V4_a1        GTTGACGCTTCTCGTTTGTCTACTCAAGGTTTCGCTTGGGATCAACCGATTGCTGACAAC  936
DSM_V4_1_a1        GTTGACGCTTCTCGTTTGTCTACTCAAGGTTTCGCTTGGGATCAACCGATTGCTGACAAC  936
```

```
BZICU-2_V5_a1     GTTGACGCTTCTCGTTTGTCTACTCAAGGTTTCGCTTGGGATCAACCGATTGCTGACAAC  948

ACICU_V1_a1       AAAACTAAAGAAGGTCGTGCTATGAACCGTCGTGTATTCGCGACAATCACTGGTAGCCGT 1017
TG02017_V1_1_a1   AAAACTAAAGAAGGTCGTGCTATGAACCGTCGTGTATTCGCGACAATCACTGGTAGCCGT 1017
AB0057_V2_a1      AAAACTAAAGAAGGTCGTGCTATGAACCGTCGTGTATTCGCGACAATCACTGGTAGCCGT 1008
4190_V3_a1        AAAACTAAAGAAGGTCGTGCTATGAACCGTCGTGTATTCGCGACAATCACTGGTAGCCGT  993
AB900_V4_a1       AAAACTAAAGAAGGTCGTGCTATGAACCGTCGTGTATTCGCGACAATCACTGGTAGCCGT  996
DSM_V4_1_a1       AAAACTAAAGAAGGTCGTGCTATGAACCGTCGTGTATTCGCGACAATCACTGGTAGCCGT  996
BZICU-2_V5_a1     AAAACTAAAGAAGGTCGTGCTATGAACCGTCGTGTATTCGCGACAATCACTGGTAGCCGT 1008

ACICU_V1_a1       ACTGTAGTAGTTCAACCTGGTCAAgAAgcggcagctcctgcagcaGCTCAATAA 1071
TG02017_V1_1_a1   ACTGTAGTAGTTCAACCTGGTCAAcAA------------------GCTCAATAA 1053
AB0057_V2_a1      ACTGTAGTAGTTCAACCTGGTCAAgAAgcggcagctcctgcagcaGCTCAATAA 1062
4190_V3_a1        ACTGTAGTAGTTCAACCTGGTCAAcAA------------------GCTCAATAA 1029
AB900_V4_a1       ACTGTAGTAGTTCAACCTGGTCAAgAAgcggcagctcctgcagcaGCTCAATAA 1050
DSM_V4_1_a1       ACTGTAGTAGTTCAACCTGGTCAAcAA------------------GCTCAATAA 1032
BZICU-2_V5_a1     ACTGTAGTAGTTCAACCTGGTCAAcAA------------------GCTCAATAA 1044
```
