## Supplementary material for "Microevolution in the major outer membrane protein OmpA of *Acinetobacter baumannii*": Figure S3

**Figure S3.** Synonymous and non-synonymous mutations among *Acinetobacter ompA* V1 and V1_1 alleles. **A.** Comparison of V1 and V1_1 nucleotide sequences. DNA alignments of the indicated 23 alleles were done using ClustalW (http://www.genome.jp/tools/clustalw/) using 1,041 nucleotide positions equivalent to 347 codons, omitting the last nucleotides of the gene encoding the variable C-terminal region. The numbers at the right indicate the position of the last nucleotide residue in the corresponding row of alignments. For short, *A. baumannii* V1 *ompA* alleles displaying the same DNA sequence in a particular row have been combined (indicated at the left). For non-*A. baumannii* sequences, GG2 indicates the *ompA* V1_1a3 allele from *A. seifertii* GG2 and OIFC021 the V1_1a4 allele from *A. nosocomialis* OIFC021 (Table S2). Nucleotide substitutions for a particular allele or groups of alleles as compared to *A. baumannii* V1a1 *ompA* are highlighted in bold red in the alignments, with the amino acid residue encoded by the triplet in which the mutations were detected (Aa involved) highlighted in bold below the corresponding sequences. It is also indicated whether the type of nucleotide substitution represented a transition (t) or a transversion (T), and whether it resulted in a synonymous (Sy) or a non-synonymous (nSy) change. OM topology predictions for the translated sequences are also indicated in the upper sequence for each set of rows. The externally exposed coding regions (EL) are highlighted in light blue, the transmembrane (TM) regions in red, and the periplasmatic domain in green. **B.** Intra- and inter-species amino acid substitutions and possible effects. The properties of the amino acids and whether the change is neutral, favored, or disfavored were from (M.J. Betts, R.B. Russell. Amino acid properties and consequences of substitutions. In Bioinformatics for Geneticists, M.R. Barnes, I.C. Gray eds, Wiley, 2003; http://www.russelllab.org/aas/). Substitutions present in only one allele (highlighted in yellow) were excluded to remove ambiguous signals from possible sequencing errors.

**A)**

```
v1a1-19           ATGAAATTGAGTCGTATTGCACTTGCTACTATGCTTGTTGCTGCTCCATTAGCTGCTGCT   60
v_1_1a1-2         ATGAAATTGAGTCGTATTGCACTTGCTACTATGCTTGTTGCTGCTCCATTAGCTGCTGCT
GG2 (v1_1a3)      ATGAAATTGAGTCGTATTGCACTTGCTACTATGCTTGTTGCTGCTCCATTAGCTGCTGCT
OIFC021 (v1_1a4)ATGAAATTGAGTCGTATTGCACTTGCTACTATGCTTGTTGCTGCTCCATTAGCTGCTGCT

                                              TM1                     EL1
v1a1-3            AATGCTGGCGTAACAGTTACTCCATTACTGCTTGGTTACACTTTCCAAGACAGCCAACAC  120
Aa involved                                 Leu
v1a4-19           AATGCTGGCGTAACAGTTACTCCATTATTGCTTGGTTACACTTTCCAAGACAGCCAACAC
v_1_1a1-2         AATGCTGGCGTAACAGTTACTCCATTATTGCTTGGTTACACTTTCCAAGACAGCCAACAC
Aa involved                                 Leu
Type of change                              t
Syn or non-Syn                              Sy

GG2               AATGCTGGCGTAACAGTTACTCCATTATTGCTTGGTTACACTTTCCAAGACAGCCAACAC  120
OIFC021           AATGCTGGCGTAACAGTTACTCCATTATTGCTTGGTTACACTTTCCAAGACAGCCAACAC
Aa involved                                 Leu
```

```
Type of change                             t
Syn or non-Syn                             Sy

                               EL1  (cont.)                             TM2
v1a1-4          AACAATGGCGGTAAAGATGGTAACTTAACTAATTCACCTGAGTTACAAGACGATTTATTC  180
Aa involved     Asn                  AsnLeu   AsnSer
v1a5,10-12,18   AACAATGGCGGTAAAGATGGTAACTTAACTAATGCACCTGAGTTACAAGACGATTTATTC
v_1_1a1-2       AACAATGGCGGTAAAGATGGTAACTTAACTAATGCACCTGAGTTACAAGACGATTTATTC
Aa involved                          AsnLeu   AsnAla
v1a16,17        AACAATGGCGGTAAAGATGGTAACTTAACTAACGGACCTGAGTTACAAGACGATTTATTC
v1a6-8,13-15,19 AACAATGGCGGTAAAGATGGTAACTTAACTAACGGTCCTGAGTTACAAGACGATTTATTC
v_1_1a2         AACAATGGCGGTAAAGATGGTAACTTAACTAACGGTCCTGAGTTACAAGACGATTTATTC
v1a9            AACAATGGCGGTAAAGATGGTAATTTAACTAACGGTCCTGAGTTACAAGACGATTTATTC
Aa involved                          AsnLeu   AsnGly
Type of change                         t        tTTT
Syn or non-Syn                        Sy      Sy nSy

GG2             AATAATGGCGGTAAAGATGGTAACCTAACTAACGGTCCTGAGTTACAAGACGATTTATTC  180
OIFC021         AATAATGGCGGTAAAGATGGTAACCTAACTAACGGTCCTGAGTTACAAGACGATTTATTC
Aa involved     Asn                     Leu   AsnGly
Type of change    t                     t       tTTT
Syn or non-Syn   Sy                     Sy    Sy nSy

                  TM2  (cont.)                IL1          TM3
v1a1-4          GTTGGTGCAGCTCTTGGTATCGAGTTAACTCCATGGTTAGGTTTCGAAGCTGAATATAAC   240
Aa involved        Gly         GlyIleGlu   Thr
v1a5,9-12,18    GTTGGCGCAGCTCTTGGTATCGAGTTAACACCATGGTTAGGTTTCGAAGCTGAATATAAC
v_1_1a1         GTTGGCGCAGCTCTTGGTATCGAGTTAACACCATGGTTAGGTTTCGAAGCTGAATATAAC
v1a6-8,13-15,19 GTTGGCGCAGCTCTTGGTATCGAGTTAACTCCATGGTTAGGTTTCGAAGCTGAATATAAC
v_1_1a2         GTTGGCGCAGCTCTTGGTATCGAGTTAACTCCATGGTTAGGTTTCGAAGCTGAATATAAC
v1a16,17        GTTGGTGCAGCTCTTGGTATCGAGTTAACACCATGGTTAGGTTTCGAAGCTGAATATAAC
Aa involved        Gly                     Thr
Type of change       t                       T
Syn or non-Syn      Sy                      Sy

GG2             GTTGGTGCAGCTCTTGGCGTTGAATTAACACCATGGTTAGGTTTCGAAGCTGAATATAAC  240
Aa involved                    GlyValGlu   Thr
Type of change                   tt t  t     T
Syn or non-Syn                 Sy nSy Sy    Sy
OIFC021         GTTGGTGCAGCTCTTGGTGTTGAATTAACACCATGGTTAGGTTTCGAAGCTGAATATAAC
Aa involved                    GlyValGlu   Thr
Type of change                    t t  t     T
Syn or non-Syn                    nSy Sy    Sy

                               EL2
v1a1-9,13-15    CAAGTTAAAGGCGACGTAGACGGCGCTTCTGCTGGTGCTGAATATAAACAAAAACAAATC   300
v_1_1a2         CAAGTTAAAGGCGACGTAGACGGCGCTTCTGCTGGTGCTGAATATAAACAAAAACAAATC
Aa involved              GlyAsp   AspGlyAlaSer            Tyr
v1a10-12,18     CAAGTTAAAGGTGACGTAGACGGCGCTTCTGCTGGTGCTGAATATAAACAAAAACAAATC
v_1_1a1         CAAGTTAAAGGTGACGTAGACGGCGCTTCTGCTGGTGCTGAATATAAACAAAAACAAATC
Aa involved              Gly
v1a16,17        CAAGTTAAAGGCGACGTAGACGGCGCTGCTGCTGGTGCTGAATATAAACAAAAACAAATC
Aa involved                                Ala
v1a19           CAAGTTAAAGGCGACGTAGACGGTCCTGTTGCTGGTGCTGAATATAAACAAAAACAAATC
Aa involved                          GlyProVal
Type of change             t           tT  Tt
Syn or non-syn            Sy         Sy nSy nSy

GG2             CAAGTTAAAGGTGATGTAAACGGCGCTTCTGCTGGTGCTGAATACAAACAAAAACAAATC  300
OIFC021         CAAGTTAAAGGTGATGTAAACGGCGCTTCTGCTGGTGCTGAATACAAACAAAAACAAATC
Aa involved              GlyAsp   Asn                     Tyr
Type of change             t  t   t                         t
Syn or non-syn           Sy  Sy   nSy                      Sy
```

```
                           TM4                      IL2             TM5
v1a1-15,18,19   AACGGTAACTTCTATGTTACTTCTGATTTAATTACTAAAAACTACGACAGCAAAATCAAG   360
v_1_1a1,2       AACGGTAACTTCTATGTTACTTCTGATTTAATTACTAAAAACTACGACAGCAAAATCAAG
Aa involved                    Val            Ile         Tyr         Ile
v1a16,17        AACGGTAACTTCTATGTTACTTCTGATTTAATCACTAAAAACTACGACAGCAAAATCAAG
Type of change                                  t
Syn or non-syn                                 Sy

GG2             AACGGTAACTTCTATGCTACTTCTGATTTAATCACTAAAAACTATGACAGCAAATTTAAG  360
Aa involved                    Ala            Ile         Tyr         Phe
Type of change                  t               t           t         T t
Syn or non-syn                 nSy             Sy          Sy         nSy
OIFC021         AACGGTAACTTCTATGCTACTTCTGATTTAATCACTAAAAACTACGACAGCAAATTTAAG
Aa involved                    Ala            Ile                     Phe
Type of change                  t               t                     T t
Syn or non-syn                 nSy             Sy                     nSy

                 TM5  (cont.)                              EL3
v1a1,2          CCGTACGTATTATTAGGTGCTGGTCACTATAAATATGACTTTGATGGCGTAAATCGTGGT   420
Aa involved     Pro      LeuLeu            TyrLysTyrAsp      Gly   Asn
v1a11,18        CCGTACGTATTATTAGGTGCTGGTCACTATAAATACGACTTTGATGGCGTAAATCGTGGT
v1a3-10,13-15   CCGTACGTATTATTAGGTGCTGGTCACTATAAATACGACTTTGATGGCGTAAACCGTGGT
v_1_1a1,2       CCGTACGTATTATTAGGTGCTGGTCACTATAAATACGACTTTGATGGCGTAAACCGTGGT
v1a12           CCGTACGTATTATTAGGTGCTGGTCACTACAAGTATGACTTTGATGGCGTAAATCGTGGT
v1a19           CCGTACGTATTATTAGGTGCTGGTCACTATAAATACGATTTTGATGGCGTAAACCGTGGT
v1a16,17        CCATACGTATTGTTAGGTGCTGGTCACTACAAGTATGACTTTGATGGCGTAAACCGTGGT
Aa involved     Pro      Leu               TyrLysTyrAsp            Asn
Type of change    t        t                 t  t  t  t             t
Syn or non-syn   Sy       Sy                Sy Sy Sy Sy            Sy

GG2             CCGTATGTATTATTGGGTGCTGGTCACTACAAATACGACTTTGATGGTGTAAATCGTGGT  420
OIFC021         CCGTATGTATTATTGGGTGCTGGTCACTACAAATACGACTTTGATGGTGTAAATCGTGGT
Aa involved                 Leu            Tyr   Tyr         Gly
Type of change                t              t     t           t
Syn or non-syn               Sy             Sy    Sy          Sy

                  EL3  (cont.)                           TM6
v1a1,2,5-9,15   ACACGTGGTACTTCTGAAGAAGGTACTTTAGGTAACGCTGGTGTTGGTGCTTTCTGGCGC   480
v_1_1a1,2       ACACGTGGTACTTCTGAAGAAGGTACTTTAGGTAACGCTGGTGTTGGTGCTTTCTGGCGC
Aa involved              ThrSer                 AsnAla    Val   Ala   Trp
v1a3,4,10-14    ACACGTGGTAACTCAGAAGAAGGTACTTTAGGTAACGCTGGTGTTGGTGCTTTCTGGCGC
v1a18,19        ACACGTGGTAATTCAGAAGAAGGTACTTTAGGTAACGCTGGTGTTGGTGCTTTCTGGCGC
v1a16,17        ACACGTGGTAACTCAGAAGAAGGTACTTTAGGTAATGCGGGTGTTGGTGCTTTCTGGCGC
Aa involved              AsnSer                 AsnAla
Nt change                 Tt   T                  t  T
Syn or non-syn           nSy  Sy                 Sy Sy

GG2             ACACGTGGTAATTCTGAAGAAGGTACTTTAGGTAATGCTGGTATCGGTGCGTTCTATCGC  480
OIFC021         ACACGTGGTAATTCTGAAGAAGGCACTTTAGGTAATGCTGGTATCGGTGCATTCTATCGC
Aa involved              Asn                    Asn      Ile   Ala   Tyr
Type of change            T                       t      t t     T    tT
Syn or non-syn           nSy                     Sy      nSy    Sy   nSy

                     IL3                TM7                            EL4
v1a1-15,17-19   TTAAACGACGCTTTATCTCTTCGTACTGAAGCTCGTGCTACTTATAATGCTGATGAAGAG  540
v_1_1a1,2       TTAAACGACGCTTTATCTCTTCGTACTGAAGCTCGTGCTACTTATAATGCTGATGAAGAG
Aa involved     LeuAsnAsp               Thr         Ala      AsnAla      Glu
v1a16           TTAAACGATGCTTTATCTCTTCGTACAGAAGCTCGTGCTACTTATAACGCTGATGAAGAG
Type of change          t                 T                    t
Syn or non-syn         Sy                Sy                   Sy

GG2             ATCAATGATGCTTTATCTCTTCGTACAGAAGCTCGTGGTACTTATAACTTTGATGAAGAA  540
Aa involved     IleAsnAsp               Thr         Gly      AsnPhe      Glu
Type of change  T T  t  t                 T           T        tTt         t
Syn or non-syn  nSy Sy Sy                Sy          nSy      Sy nSy      Sy
OIFC021         ATCAACGATGCTTTATCTCTTCGTACAGAAGCTCGTGGTACTTATAACTTTGATGAACAA
Aa involved     Ile   Asp               Thr         Gly      AsnPhe      Gln
Type of change  T T     t                 T           T        tTt       T t
```

```
Syn or non-syn  nSy    Sy                Sy         nSy      Sy nSy      nSy

                EL4(cont.)              TM8                periplasmic domain
v1a1,3-15,17-19 TTCTGGAACTATACAGCTCTTGCTGGCTTAAACGTAGTTCTTGGTGGTCACTTGAAGCCT   600
v_1_1a1,2       TTCTGGAACTATACAGCTCTTGCTGGCTTAAACGTAGTTCTTGGTGGTCACTTGAAGCCT
Aa involved                          AlaGly
v1a2            TTCTGGAACTATACAGCTCTTGCAGGCTTAAACGTAGTTCTTGGTGGTCACTTGAAGCCT
Aa involved                          Ala
Type of change                         T
Syn or non-syn                        Sy
v1a16           TTCTGGAACTATACAGCTCTTGCTGGTTTAAACGTAGTTCTTGGTGGTCACTTGAAGCCT
Aa involved                             Gly
Type of change                            t
Syn or non-syn                           Sy

GG2             TTCTGGAACTATACAGCTCTTGCTGGTTTAAACGTAGTTCTTGGTGGTCACTTGAAGCCT  600
OIFC021         TTCTGGAACTATACAGCTCTTGCTGGTTTAAACGTAGTTCTTGGTGGTCACTTGAAGCCT
Aa involved                             Gly
Type of change                            t
Syn or non-syn                           Sy

                                         periplasmic domain (cont.)
v1a1-14,16-19   GCTGCTCCTGTAGTAGAAGTTGCTCCAGTTGAACCAACTCCAGTTGCTCCACAACCACAA  660
v_1_1a1-2       GCTGCTCCTGTAGTAGAAGTTGCTCCAGTTGAACCAACTCCAGTTGCTCCACAACCACAA
Aa involved                                                  Ala
v1a15           GCTGCTCCTGTAGTAGAAGTTGCTCCAGTTGAACCAACTCCAGTTACTCCACAACCACAA
Aa involved                                                  Thr
Type of change                                               t
Syn or non-syn                                               nSy

GG2             GCTGCTCCTGTAGTAGAAGTTGCTCCAGTTGAACCAACTCCAGTTGCTCCACAACCACAA  660
OIFC021         GCTGCTCCTGTAGTAGAAGTTGCTCCAGTTGAACCAACTCCAGTTGCTCCACAACCACAA



                                         periplasmic domain (cont.)
v1a1-19         GAGTTAACTGAAGACCTTAACATGGAACTTCGTGTGTTCTTTGATACTAACAAATCAAAC  720
v_1_1a1-2       GAGTTAACTGAAGACCTTAACATGGAACTTCGTGTGTTCTTTGATACTAACAAATCAAAC
Aa involved                                                        Lys

OIFC021         GAGTTAACTGAAGACCTTAACATGGAACTTCGTGTGTTCTTTGATACTAACAAATCAAAC  720
GG2             GAGTTAACTGAAGACCTTAACATGGAACTTCGTGTGTTCTTTGATACTAACAAGTCAAAC
Aa involved                                                        Lys
Type of change                                                       t
Syn or non-syn                                                      Sy

                                         periplasmic domain (cont.)
v1a1,2,6,10,12  ATCAAAGACCAATACAAGCCAGAAATCGCTAAAGTTGCTGAAAAATTATCTGAATACCCT  780
v1a16           ATCAAAGACCAATACAAGCCAGAAATCGCTAAAGTTGCTGAAAAATTATCTGAATACCCT
v_1_1a1,2       ATCAAAGACCAATACAAGCCAGAAATCGCTAAAGTTGCTGAAAAATTATCTGAATACCCT
Aa involved                             Ile            GluLys   Ser
v1a3-5,7-9      ATCAAAGACCAATACAAGCCAGAAATTGCTAAAGTTGCTGAAAAATTATCTGAATACCCT
v1a11,13-19     ATCAAAGACCAATACAAGCCAGAAATTGCTAAAGTTGCTGAAAAATTATCTGAATACCCT
Aa involved                             Ile
Type of change                            t
Syn or non-syn                           Sy

OIFC021         ATCAAAGACCAATACAAGCCAGAAATCGCTAAAGTTGCTGAAAAATTATCTGAATACCCT
GG2             ATCAAAGACCAATACAAGCCAGAAATCGCTAAAGTTGCTGAGAAGTTAACTGAATACCCT  780
Aa involved                                            GluLys   Thr
Type of change                                           t  t   T
Syn or non-syn                                          Sy Sy   nSy

                                         periplasmic domain (cont.)
v1a1-13,15-19   AACGCTACTGCACGTATCGAAGGTCACACAGATAACACTGGTCCACGTAAGTTGAACGAA  840
```

```
v_1_1a1,2       AACGCTACTGCACGTATCGAAGGTCACACAGATAACACTGGTCCACGTAAGTTGAACGAA
v1a14           AACGCTACTGCACGTATCGAAGGCCACACAGATAACACTGGTCCACGTAAGTTGAACGAA
Aa involved                          Gly
Type of change                         t
Syn or non-syn                        Sy

GG2             AACGCTACTGCACGTATCGAAGGTCACACAGATAACACTGGTCCACGTAAGTTGAACGAA  840
OIFC021         AACGCTACTGCACGTATCGAAGGTCACACAGATAACACTGGTCCACGTAAGTTGAACGAA

                                         periplasmic domain (cont.)
v1a1-19         CGTTTATCTTTAGCTCGTGCTAACTCTGTTAAATCAGCTCTTGTAAACGAATACAACGTT  900
Aa involved                                                        TyrAsn
v_1_1a1,2       CGTTTATCTTTAGCTCGTGCTAACTCTGTTAAATCAGCTCTTGTAAACGAATATAACGTT
Aa involved                                                        Tyr
Type of change                                                       t
Syn or non-syn                                                      Sy

GG2             CGTTTATCTTTAGCTCGTGCTAACTCTGTTAAATCAGCTCTTGTAAACGAATACAATGTT  900
OIFC021         CGTTTATCTTTAGCTCGTGCTAACTCTGTTAAATCAGCTCTTGTAAACGAATACAATGTT
Aa involved                                                           Asn
Type of change                                                          t
Syn or non-syn                                                         Sy

                                         periplasmic domain (cont.)
v1a1-4,6,7,16   GACGCTTCTCGTTTGTCTACTCAAGGTTTCGCTTGGGATCAACCGATTGCTGACAACAAA  960
v1a10-12        GACGCTTCTCGTTTGTCTACTCAAGGTTTCGCTTGGGATCAACCGATTGCTGACAACAAA
v_1_1a1,2       GACGCTTCTCGTTTGTCTACTCAAGGTTTCGCTTGGGATCAACCGATTGCTGACAACAAA
Aa change       Asp
v1a5,8,9,13-15  GATGCTTCTCGTTTGTCTACTCAAGGTTTCGCTTGGGATCAACCGATTGCTGACAACAAA
v1a17-19        GATGCTTCTCGTTTGTCTACTCAAGGTTTCGCTTGGGATCAACCGATTGCTGACAACAAA
Aa change       Asp
Type of change    t
Syn or non-syn   Sy

OIFC021         GACGCTTCTCGTTTGTCTACTCAAGGTTTCGCTTGGGATCAACCGATTGCTGACAACAAA  960
GG2             GATGCTTCTCGTTTGTCTACTCAAGGTTTCGCTTGGGATCAACCGATTGCTGACAACAAA
Aa change       Asp
Type of change    t
Syn or non-syn   Sy

                                         periplasmic domain (cont.)
v1a1-19         ACTAAAGAAGGTCGTGCTATGAACCGTCGTGTATTCGCGACAATCACTGGTAGCCGTACT 1020
v_1_1a1,2       ACTAAAGAAGGTCGTGCTATGAACCGTCGTGTATTCGCGACAATCACTGGTAGCCGTACT
GG2             ACTAAAGAAGGTCGTGCTATGAACCGTCGTGTATTCGCGACAATCACTGGTAGCCGTACT
OIFC021         ACTAAAGAAGGTCGTGCTATGAACCGTCGTGTATTCGCGACAATTACTGGTAGCCGTACT

v1a1-19         GTAGTAGTTCAACCTGGTCAA 1041
v_1_1a1,2       GTAGTAGTTCAACCTGGTCAA
GG2             GTAGTAGTTCAACCTGGTCAA
OIFC021         GTAGTAGTTCAACCTGGTCAA
```

**B) Amino acid substitutions and possible effects**

**At EL1:**

Among *A. baumannii* alleles:

Ser52 (4 alleles including V1a1): polar side chain, uncharged, contains hydroxyl group

Ala52 (6 alleles): hydrophobic side chain

Gly52 (11 alleles): minimal side chain having one hydrogen atom, considered polar.

In non-*baumannii* alleles (GG2/OIFC021)

Gly52 in both cases

In all cases the changes are described as conservative (neutral or slightly favored).

**At EL2:**
Among *A. baumannii* alleles
Ala89 (20 alleles): hydrophobic side chain
Pro89 (Present in only one allele: V1a19).
In non-*baumannii* alleles (GG2/OIFC021)
GG2/OIFC021: Ala89

Among *A. baumannii* alleles
Ser90 (18 alleles, including V1a1): polar side chain, uncharged, contains hydroxyl group
Ala90 (2 alleles: v1a16,17): hydrophobic side chain
Val90 (Present in only one allele: V1a19).
In non-*baumannii* alleles (GG2/OIFC021)
GG2/OIFC021: Ser90
Intra-species amino acid change, described as favored

Asp87 (all 21 *A. baumannii* alleles):, polar side chain, charged, contains carboxyl group (acidic).
In non-*baumannii* alleles
GG2/OIFC021: Asn87: amide of aspartic acid, polar side chain, uncharged, high propensity to hydrogen bond.
Inter-species change, described as favored.

**At EL3:**
Among *A. baumannii* alleles
Thr144 (10 alleles, including V1a1): polar, uncharged, contains hydroxyl group
Asn144 (11 alleles): polar, uncharged, amide of aspartic acid.
Intra-species amino acid change, described as neutral.
In non-*baumannii* alleles
In GG2/OIFC021: Asn144

**At EL4:**
Ala177 (all 21 *A. baumannii* alleles): hydrophobic.
In non-*baumannii* alleles
GG2/OIFC021: Phe177: aromatic side chain, highly hydrophobic, bulky.
Inter-species amino acid change, described as ***disfavored***.

Glu180 (all 21 *A. baumannii* alleles): polar, charged, contains carboxyl group (acidic).
In non-*baumannii* alleles
GG2/OIFC021: Gln180: polar, uncharged. High propensity to hydrogen bond.
Inter-species amino acid changes, described as favored.

**Non-exposed regions**
**At TM4:**
Val106 (all 21 *A. baumannii* alleles): hydrophobic, binding/recognition of hydrophobic ligands such as lipids.
In non-*baumannii* alleles
GG2/OIFC021: Ala106: aliphatic side chain, highly hydrophobic.
Inter-species amino acid changes, described as neutral for membrane proteins.

**At TM5**
Ile119 (all 21 *A. baumannii* alleles): hydrophobic side chain
In non-*baumannii* alleles
GG2/OIFC021: Phe119: aromatic, highly hydrophobic, bulky.
Inter-species change, described as ***disfavored*** for membrane proteins.

**At TM6**
Val155 (All 21 *A. baumannii* alleles)
In non-*baumannii* alleles
GG2/OIFC021: Ile155
Inter-species amino acid changes, described as favored for membrane proteins.

Trp159 (All 21 *A. baumannii* alleles)
In non-*baumannii* alleles
GG2/OIFC021: Tyr159
Inter-species amino acid changes, described as ***disfavored*** for membrane proteins.

In A. *baumannii*
Leu161 (all 21 *A. baumannii* alelles): hydrophobic
In non-*baumannii* alleles
GG2/OIFC021: Ile161: hydrophobic.
Inter-species amino acid changes, described as favored for membrane proteins.

**At TM7**
Ala173 (all 21 *A. baumannii* alleles): hydrophobic.
In non-*baumannii* alleles
GG2/OIFC021: Gly173.
Inter-species amino acid change, described as favored for membrane proteins.

**At the periplasmic domain**
Ala216 (20 *A. baumannii* alelles): hydrophobic.
Thr216. Present in only one allele: V1a15:
In non-*baumannii* alleles
GG2/OIFC021: Ala216.
