## Supplementary material for "Microevolution in the major outer membrane protein OmpA of *Acinetobacter baumannii*": Table S1

**Table S1.** ***ompA*** **variant alleles and ST profiles (Pasteur scheme) of the** ***A. baumannii*** **strains analyzed in this work.**

| Variant/allele | Strain | housekeeping gene alleles[a] | | | | | | | | |
|---|---|---|---|---|---|---|---|---|---|---|
| | | **cpn60** | **fusA** | **gltA** | **pyrG** | **recA** | **rplB** | **rpoB** | **ST[a]** | **CC[b]** |
| **V1 group** | | | | | | | | | | |
| V1a1 | PKAB07 | 2 | nd | 2 | 2 | 2 | 2 | 2 | nd[c] | 2* |
| | 48055 | 2 | 2 | 2 | 2 | 2 | 2 | 2 | 2 | 2 |
| | 48055 | 2 | 2 | 2 | 2 | 2 | 2 | 2 | 2 | 2 |
| | 53264 | 2 | 2 | 2 | 2 | 2 | 2 | 2 | 2 | 2 |
| | Ab11111 | 2 | 2 | 2 | 2 | 2 | 2 | 2 | 2 | 2 |
| | AB1H8 | 2 | 2 | 2 | 2 | 2 | 2 | 2 | 2 | 2 |
| | AB_1582-8 | 2 | 2 | 2 | 2 | 2 | 2 | 2 | 2 | 2 |
| | AB_1595-8 | 2 | 2 | 2 | 2 | 2 | 2 | 2 | 2 | 2 |
| | AB_1766_8 | 2 | 2 | 2 | 2 | 2 | 2 | 2 | 2 | 2 |
| | AB_2008-23-07-0 | 2 | 2 | 2 | 2 | 2 | 2 | 2 | 2 | 2 |
| | AB_2009-04-02 | 2 | 2 | 2 | 2 | 2 | 2 | 2 | 2 | 2 |
| | AB_515-8 | 2 | 2 | 2 | 2 | 2 | 2 | 2 | 2 | 2 |
| | AB_908-12 | 2 | 2 | 2 | 2 | 2 | 2 | 2 | 2 | 2 |
| | AB_909-05 | 2 | 2 | 2 | 2 | 2 | 2 | 2 | 2 | 2 |
| | AB_TG2631 | 2 | 2 | 2 | 2 | 2 | 2 | 2 | 2 | 2 |
| | AB_TG27323 | 2 | 2 | 2 | 2 | 2 | 2 | 2 | 2 | 2 |
| | AB_TG27327 | 2 | 2 | 2 | 2 | 2 | 2 | 2 | 2 | 2 |
| | AB_TG27331 | 2 | 2 | 2 | 2 | 2 | 2 | 2 | 2 | 2 |
| | AB_TG27335 | 2 | 2 | 2 | 2 | 2 | 2 | 2 | 2 | 2 |
| | LY4 | 2 | 2 | na | 2 | 2 | 2 | 2 | 2 | 2 |
| | Naval-113 | 2 | 2 | 2 | 2 | 2 | 2 | 2 | 2 | 2 |
| | NIPH 2061 | 2 | 2 | 2 | 2 | 2 | 2 | 2 | 2 | 2 |
| | TG2013 | 2 | 2 | 2 | 2 | 2 | 2 | 2 | 2 | 2 |
| | TG2014 | 2 | 2 | 2 | 2 | 2 | 2 | 2 | 2 | 2 |
| | TG22202 | 2 | 2 | 2 | 2 | 2 | 2 | 2 | 2 | 2 |
| | TG27299 | 2 | 2 | 2 | 2 | 2 | 2 | 2 | 2 | 2 |
| | TG27315 | 2 | 2 | 2 | 2 | 2 | 2 | 2 | 2 | 2 |
| | TG27319 | 2 | 2 | 2 | 2 | 2 | 2 | 2 | 2 | 2 |
| | TG27371 | 2 | 2 | 2 | 2 | 2 | 2 | 2 | 2 | 2 |
| | TG27379 | 2 | 2 | 2 | 2 | 2 | 2 | 2 | 2 | 2 |
| | TG27383 | 2 | 2 | 2 | 2 | 2 | 2 | 2 | 2 | 2 |
| | TG27407 | 2 | 2 | 2 | 2 | 2 | 2 | 2 | 2 | 2 |
| | TG27411 | 2 | 2 | 2 | 2 | 2 | 2 | 2 | 2 | 2 |
| | AB210 | 2 | 2 | 2 | 2 | 2 | 2 | 2 | 2 | 2 |
| | BJAB07104 | 2 | 2 | 2 | 2 | 2 | 2 | 2 | 2 | 2 |
| | TCDC-AB0715 | 2 | 2 | 2 | 2 | 2 | 2 | 2 | 2 | 2 |
| | W7282 | 2 | 2 | 2 | 2 | 2 | 2 | 2 | 2 | 2 |
| | ABIsac_ColiR | 2 | 2 | 2 | 2 | 2 | 2 | 2 | 2 | 2 |
| | ABIsac_ColiS | 2 | 2 | 2 | 2 | 2 | 2 | 2 | 2 | 2 |
| | Ab44444 | 2 | 2 | 2 | 2 | 2 | 2 | 2 | 2 | 2 |
| | ABNIH1 | 2 | 2 | 2 | 2 | 2 | 2 | 2 | 2 | 2 |
| | ABNIH18 | 2 | 2 | 2 | 2 | 2 | 2 | 2 | 2 | 2 |
| | ABNIH25 | 2 | 2 | 2 | 2 | 2 | 2 | 2 | 2 | 2 |
| | ABNIH5 | 2 | 2 | 2 | 2 | 2 | 2 | 2 | 2 | 2 |
| | Naval-2 | 2 | 2 | 2 | 2 | 2 | 2 | 2 | 2 | 2 |
| | Naval-78 | 2 | 2 | 2 | 2 | 2 | 2 | 2 | 2 | 2 |
| | TG15241 | 2 | 2 | 2 | 2 | 2 | 2 | 2 | 2 | 2 |
| | TG15242 | 2 | 2 | 2 | 2 | 2 | 2 | 2 | 2 | 2 |
| | TG20546 | 2 | 2 | 2 | 2 | 2 | 2 | 2 | 2 | 2 |
| | ABNIH13 | 2 | 2 | 2 | 2 | 2 | 2 | 2 | 2 | 2 |
| | ABNIH14 | 2 | 2 | 2 | 2 | 2 | 2 | 2 | 2 | 2 |
| | ABNIH15 | 2 | 2 | 2 | 2 | 2 | 2 | 2 | 2 | 2 |
| | ABNIH16 | 2 | 2 | 2 | 2 | 2 | 2 | 2 | 2 | 2 |
| | ABNIH17 | 2 | 2 | 2 | 2 | 2 | 2 | 2 | 2 | 2 |
| | ABNIH2 | 2 | 2 | 2 | 2 | 2 | 2 | 2 | 2 | 2 |
| | ABNIH20 | 2 | 2 | 2 | 2 | 2 | 2 | 2 | 2 | 2 |
| | ABNIH22 | 2 | 2 | 2 | 2 | 2 | 2 | 2 | 2 | 2 |
| | ABNIH23 | 2 | 2 | 2 | 2 | 2 | 2 | 2 | 2 | 2 |
| | ABNIH24 | 2 | 2 | 2 | 2 | 2 | 2 | 2 | 2 | 2 |
| | ABNIH26 | 2 | 2 | 2 | 2 | 2 | 2 | 2 | 2 | 2 |
| | TG15233 | 2 | 2 | 2 | 2 | 2 | 2 | 2 | 2 | 2 |
| | TG15236 | 2 | 2 | 2 | 2 | 2 | 2 | 2 | 2 | 2 |
| | TG15238 | 2 | 2 | 2 | 2 | 2 | 2 | 2 | 2 | 2 |
| | TG15239 | 2 | 2 | 2 | 2 | 2 | 2 | 2 | 2 | 2 |
| | AB_2008-15-34 | 2 | 2 | 2 | 2 | 2 | 2 | 2 | 2 | 2 |
| | TG15234 | 2 | 2 | 2 | 2 | 2 | 2 | 2 | 2 | 2 |
| | UMB001 | 2 | 2 | 2 | 2 | 2 | 2 | 2 | 2 | 2 |
| | NIPH 528 | 2 | 2 | 2 | 2 | 2 | 2 | 2 | 2 | 2 |
| | OIFC338 | 2 | 2 | 2 | 2 | 2 | 2 | 2 | 2 | 2 |
| | BJAB0868 | 2 | 2 | 2 | 2 | 2 | 2 | 2 | 2 | 2 |
| | ZWS1122 | 2 | 2 | 2 | 2 | 2 | 2 | 2 | 2 | 2 |
| | ZWS1219 | 2 | 2 | 2 | 2 | 2 | 2 | 2 | 2 | 2 |
| | MDR-TJ | 2 | 2 | 2 | 2 | 2 | 2 | 2 | 2 | 2 |
| | 1656-2 | 2 | 2 | 2 | 2 | 2 | 2 | 2 | 2 | 2 |
| | AB_908-14-7 | 2 | 2 | 2 | 2 | 2 | 2 | 2 | 2 | 2 |
| | DU202 | 2 | 2 | 2 | 2 | 2 | 2 | 2 | 2 | 2 |
| | 3990 | 2 | 2 | 2 | 2 | 2 | 2 | 2 | 2 | 2 |
| | ACICU | 2 | 2 | 2 | 2 | 2 | 2 | 2 | 2 | 2 |
| | TG15240 | 2 | 2 | 2 | 2 | 2 | 2 | 2 | 2 | 2 |
| | 5711 | 2 | 2 | 2 | 2 | 2 | 2 | 2 | 2 | 2 |
| | TYTH-1 | 2 | 2 | 2 | 2 | 2 | 2 | 2 | 2 | 2 |
| | LY7 | 2 | 2 | 2 | 2 | 2 | 2 | 2 | 2 | 2 |
| | NIPH24 | 2 | 2 | 2 | 2 | 2 | 2 | 2 | 2 | 2 |
| | MDR-ZJ06 | 2 | 2 | 2 | 2 | 2 | 2 | 2 | 2 | 2 |
| | OIFC180 | 2 | 2 | 2 | 2 | 2 | 2 | 2 | 2 | 2 |
| | 6014059 | 2 | 2 | 2 | 2 | 2 | 2 | 2 | 2 | 2 |
| | AB_TG5064 | 2 | 2 | 2 | 2 | 2 | 2 | 2 | 2 | 2 |
| | TG22212 | 2 | 2 | 2 | 2 | 2 | 2 | 2 | 2 | 2 |
| | TG15237 | 2 | 2 | 2 | 2 | 2 | 2 | 2 | 2 | 2 |

| | | | | | | | | | | |
|---|---|---|---|---|---|---|---|---|---|---|
| | AB_TG2022 | 2 | 2 | 2 | 2 | 2 | 2 | 2 | 2 | 2 |
| | AB_TG2023 | 2 | 2 | 2 | 2 | 2 | 2 | 2 | 2 | 2 |
| | TG2012 | 2 | 2 | 2 | 2 | 2 | 2 | 2 | 2 | 2 |
| | TG22110 | 2 | 2 | 2 | 2 | 2 | 2 | 2 | 2 | 2 |
| | TG22192 | 2 | 2 | 2 | 2 | 2 | 2 | 2 | 2 | 2 |
| | TG27307 | 2 | 2 | 2 | 2 | 2 | 2 | 2 | 2 | 2 |
| | TG27311 | 2 | 2 | 2 | 2 | 2 | 2 | 2 | 2 | 2 |
| | IS-143 | 2 | 2 | 2 | 2 | 2 | 37 | 2 | 414 | 2* |
| | NIPH1362 | 2 | 13 | 2 | 2 | 2 | 2 | 2 | 47 | 2* |
| | AB_2008-15-4! | 2 | 2 | 2 | 2 | 68 | 2 | 2 | 415 | 2* |
| | AB_2008-15-7( | 2 | 2 | 2 | 2 | 68 | 2 | 2 | 415 | 2* |
| | AB_2007-09-110-( | 3 | 74 | 2 | 3 | 6 | 1 | 16 | 429 | 1 allelic difference to ST428, see below |
| | NIPH70 | 1 | 2 | 2 | 2 | 3 | 1 | 2 | 36 | |
| V1a2 | ABNIH3 | 2 | 2 | 2 | 2 | 68 | 2 | 2 | 415 | 2* |
| V1a3 | TG02011 | 26 | 72 | 2 | 2 | 29 | 4 | 5 | 422 | |
| | AB_TG27343 | 26 | 72 | 2 | 2 | 29 | 4 | 5 | 422 | |
| | AB_1583-8 | 26 | 72 | 2 | 2 | 29 | 4 | 5 | 422 | |
| V1a4 | TG19617 | 3 | 2 | 2 | 7 | 9 | 4 | 5 | 438 | |
| | OIFC0162 | 1 | 52 | 2 | 2 | 67 | 4 | 5 | 412 | |
| | TG27387 | 5 | 4 | 4 | 1 | 3 | 3 | 4 | 6 | |
| | NCTC10304 | na | na | na | na | na | na | na | | |
| V1a5 | Ab244 (Arg) | 6 | 6 | 8 | 2 | 3 | 5 | 4 | 15 | |
| | NIPH1734 | 6 | 6 | 8 | 2 | 3 | 5 | 4 | 15 | |
| V1a6 | Ab33333 | 3 | 71 | 2 | 2 | 5 | 4 | 14 | 419 | |
| | AB4A3 | 3 | 71 | 2 | 2 | 5 | 4 | 14 | 419 | |
| | NIPH 67 | 3 | 71 | 2 | 2 | 5 | 4 | 14 | 419 | |
| V1a7 | OIFC099 | 1 | 1 | 2 | 2 | 3 | 4 | 4 | 32 | 32 |
| | OIFC087 | 1 | 1 | 2 | 2 | 3 | 4 | 4 | 32 | 32 |
| V1a8 | OIFC074 | 1 | 2 | 1 | 1 | 5 | 1 | 1 | 19 | 1* (L2)[c] |
| | Naval-21 | 1 | 2 | 1 | 1 | 5 | 1 | 1 | 19 | 1* (L2)[c] |
| | OIFC047 | 1 | 75 | 2 | 2 | 5 | 1 | 2 | 430 | |
| | ATCC19606 | 3 | 2 | 2 | 7 | 9 | 1 | 5 | 52 | |
| | JCM6841 | 3 | 2 | 2 | 7 | 9 | 1 | 5 | 52 | |
| | MRSN 3405 | 1 | 2 | 2 | 1 | 5 | 1 | 1 | 94 | 1**(L2)[c] |
| V1a9 | TG22146 | 25 | 3 | 6 | 2 | 28 | 1 | 29 | 78 | |
| | TG22144 | 25 | 3 | 6 | 2 | 28 | 1 | 29 | 78 | |
| | TG22150 | 25 | 3 | 6 | 2 | 28 | 1 | 29 | 78 | |
| V1a10 | OIFC035 | 3 | 2 | 6 | 1 | 3 | 4 | 59 | 403 | |
| | NIPH 329 | 1 | 2 | 6 | 2 | 3 | 4 | 4 | 11 | |
| V1a11 | ZW85-1 | 3 | 3 | 2 | 2 | 11 | 57 | 4 | 639 | |
| V1a12 | Naval-57 | 3 | 2 | 2 | 2 | 44 | 4 | 4 | 155 | |
| V1a13 | OIFC098 | 1 | 3 | 2 | 1 | 4 | 4 | 4 | 10 | 10 |
| | NIPH 335 | 1 | 3 | 2 | 1 | 4 | 4 | 4 | 10 | 10 |
| | BJAB0715 | 1 | 3 | 10 | 1 | 4 | 4 | 4 | 23 | 10* |
| V1a14 | AA-014 | 41 | 42 | 13 | 1 | 5 | 4 | 14 | 158 | |
| V1a15 | ATCC17978 | 3 | 2 | 2 | 2 | 30 | 4 | 28 | 437 | |
| V1a16 | D1279779 | 12 | 37 | 2 | 2 | 3 | 2 | 14 | 267 | |
| V1a17 | NIPH 410 | 10 | 4 | 3 | 2 | 13 | 1 | 2 | 39 | |
| | AB405E4 | 3 | 4 | 2 | 2 | 9 | 1 | 2 | 516 | |
| V1a18 | NIPH 615 | 3 | 4 | 2 | 2 | 9 | 1 | 2 | 516 | |
| V1a19 | TG22204 | 3 | 3 | 63 | 6 | 4 | 9 | 1 | 425 | |
| V1_1a1 | TG02017 | 3 | 1 | 7 | 1 | 7 | 1 | 4 | 424 | |
| V1_1a2 | NIPH190 | 3 | 1 | 5 | 3 | 6 | 1 | 3 | 9 | 2 allelic differences to ST428 and ST429, see above |
| | Naval-82 | 3 | 1 | 2 | 3 | 6 | 1 | 16 | 428 | 1 allelic difference to ST429, see above |
| | TG27391 | 13 | 3 | 2 | 3 | 6 | 1 | 16 | 427 | 2 allelic differences to ST428 and ST429, see above |
| **V2 group** | | | | | | | | | | |
| V2a1 | AB0057 | 1 | 1 | 1 | 1 | 5 | 1 | 1 | 1 | 1(L1)[c] |
| | AB_908-13 | 1 | 1 | 1 | 1 | 5 | 1 | 1 | 1 | 1(L1)[c] |
| | AB_909-02-7 | 1 | 1 | 1 | 1 | 5 | 1 | 1 | 1 | 1(L1)c |
| | TG22112 | 1 | 1 | 1 | 1 | 5 | 1 | 1 | 1 | 1 |
| | TG22148 | 1 | 1 | 1 | 1 | 5 | 1 | 1 | 1 | 1 |
| | TG22190 | 1 | 1 | 1 | 1 | 5 | 1 | 1 | 1 | 1 |
| | TG22194 | 1 | 1 | 1 | 1 | 5 | 1 | 1 | 1 | 1 |
| | TG22196 | 1 | 1 | 1 | 1 | 5 | 1 | 1 | 1 | 1 |
| | TG22214 | 1 | 1 | 1 | 1 | 5 | 1 | 1 | 1 | 1 |
| | AB307-0294 | 1 | 1 | 1 | 1 | 5 | 1 | 1 | 1 | 1 |
| | AB_NIH6 | 1 | 1 | 1 | 1 | 5 | 1 | 1 | 1 | 1 |
| | AB_NIH7 | 1 | 1 | 1 | 1 | 5 | 1 | 1 | 1 | 1 |
| | AB_NIH11 | 1 | 1 | 1 | 1 | 5 | 1 | 1 | 1 | 1 |
| | AB_NIH19 | 1 | 1 | 1 | 1 | 5 | 1 | 1 | 1 | 1 |
| | AYE | 1 | 1 | 1 | 1 | 5 | 1 | 1 | 1 | 1(L1)[c] |
| | MRSN 57 | 1 | 1 | 1 | 1 | 5 | 1 | 1 | 1 | 1 |
| | MRSN 58 | 1 | 1 | 1 | 1 | 5 | 1 | 1 | 1 | 1 |
| | NIPH 290 | 1 | 1 | 1 | 1 | 5 | 1 | 1 | 1 | 1(L1)[c] |
| | NIPH 527 | 1 | 1 | 1 | 1 | 5 | 1 | 1 | 1 | 1(L1)[c] |
| | TG19582 | 1 | 1 | 1 | 1 | 5 | 1 | 1 | 1 | 1 |
| | ANC4097 | 1 | 1 | 1 | 1 | 5 | 1 | 1 | 1 | 1(L1)[c] |
| | AB5075 | 1 | 1 | 1 | 1 | 5 | 1 | 1 | 1 | 1(L1)[c] |
| | IS-235 | 1 | 1 | 1 | 1 | 5 | 1 | 1 | 1 | 1 |
| | IS-251 | 1 | 1 | 1 | 1 | 5 | 1 | 1 | 1 | 1 |
| | IS-58 | 1 | 1 | 1 | 1 | 5 | 1 | 1 | 1 | 1(L1)[c] |
| | Canada BC 1 | 1 | 1 | 1 | 1 | 5 | 1 | 1 | 1 | 1(L1)[c] |
| | TG20277 | 1 | 1 | 1 | 1 | 5 | 1 | 1 | 1 | 1 |
| | Naval-83 | 3 | 1 | 1 | 1 | 5 | 1 | 1 | 20 | 1** (Two allelic differences to ST1)(L1)[c] |
| | AB_2008-15-52 | 1 | 2 | 2 | 2 | 4 | 1 | 4 | 416 | 417** (Two allelic differences to ST417) |
| | AB_2008-15-71 | 1 | 2 | 2 | 2 | 4 | 1 | 4 | 416 | 417** (Two allelic differences to ST417) |
| | AB_2008-23-01-0 | 1 | 2 | 2 | 2 | 11 | 1 | 5 | 417 | |
| | AB_2009-04-01 | 1 | 2 | 2 | 2 | 11 | 1 | 5 | 417 | |
| | AB_909-01-7 | 1 | 2 | 2 | 2 | 11 | 1 | 5 | 417 | |
| | AB_1594-8 | 1 | 2 | 2 | 2 | 11 | 1 | 5 | 417 | |
| | AB_TG2018 | 1 | 2 | 2 | 2 | 11 | 1 | 5 | 417 | |

| | | | | | | | | | | |
|---|---|---|---|---|---|---|---|---|---|---|
| | TG27395 | 1 | 2 | 2 | 2 | 11 | 1 | 5 | 417 | |
| | TG27399 | 1 | 2 | 2 | 2 | 11 | 1 | 5 | 417 | |
| | AB_1536-8 | 1 | 3 | 2 | 2 | 5 | 8 | 12 | 413 | |
| | NIPH 1669 | 3 | 3 | 2 | 2 | 3 | 1 | 3 | 3 | 3 |
| | AB4857 | 3 | 3 | 2 | 2 | 3 | 1 | 3 | 3 | 3 |
| | WC-A-694 | 3 | 3 | 2 | 2 | 3 | 1 | 3 | 3 | 3 |
| | Naval-13 | 3 | 3 | 2 | 2 | 3 | 1 | 3 | 3 | 3 |
| | WC-692 | 56 | 3 | 55 | 2 | 9 | 4 | 14 | 513 | |
| V2a2 | AB_TG2028 | 1 | 1 | 1 | 2 | 65 | 1 | 5 | 406 | |
| | AB_TG2030 | 1 | 1 | 1 | 2 | 65 | 1 | 5 | 406 | |
| | AB_TG2031 | 1 | 1 | 1 | 2 | 65 | 1 | 5 | 406 | |
| | AB_TG2032 | 1 | 1 | 1 | 2 | 65 | 1 | 5 | 406 | |
| | TG07725 | 1 | 1 | 1 | 2 | 65 | 1 | 5 | 406 | |
| | TG22332 | 1 | 1 | 1 | 2 | 65 | 1 | 5 | 406 | |
| | TG22336 | 1 | 1 | 1 | 2 | 65 | 1 | 5 | 406 | |
| | TG27295 | 1 | 1 | 1 | 2 | 65 | 1 | 5 | 406 | |
| V2a3 | OIFC065 | 3 | 2 | 19 | 25 | 5 | 2 | 5 | 136 | |
| | IS-116 | 3 | 2 | 19 | 25 | 5 | 2 | 5 | 136 | |
| V2a4 | TG00314 | 64 | 52 | 7 | 2 | 29 | 4 | 29 | 411 | |
| V2a5 | OIFC110 | 56 | 3 | 2 | 2 | 9 | 4 | 14 | 515 | |
| V2a6 | 6013113 | 1 | 1 | 1 | 1 | 5 | 1 | 2 | 81 | 1*(L2)[c] |
| | 6013150 | 1 | 1 | 1 | 1 | 5 | 1 | 2 | 81 | 1*(L2)[c] |
| **V3 group** | | | | | | | | | | |
| V3a1 | AB5256 | 3 | 3 | 2 | 4 | 7 | 2 | 4 | 25 | 25 |
| | UMB003 | 3 | 3 | 2 | 4 | 7 | 2 | 4 | 25 | 25 |
| | AB_2008-15-6 | 3 | 3 | 2 | 4 | 7 | 2 | 4 | 25 | 25 |
| | NIPH 146 | 3 | 3 | 2 | 4 | 7 | 2 | 4 | 25 | 25 |
| | 4190 | 3 | 3 | 2 | 4 | 7 | 2 | 4 | 25 | 25 |
| | AB_1649-8 | 3 | 3 | 3 | 4 | 7 | 4 | 4 | 113 | 25** |
| | AB_1650-8 | 3 | 3 | 3 | 4 | 7 | 4 | 4 | 113 | 25** |
| | WC-348 | 1 | 52 | 2 | 2 | 67 | 4 | 5 | 412 | |
| | EGD-HP18 | 3 | 1 | 2 | 1 | 18 | 1 | 48 | 221 | |
| **V4 group** | | | | | | | | | | |
| V4a1 | AB900 | 3 | 3 | 6 | 2 | 3 | 1 | 5 | 49 | |
| | OIFC111 | 3 | 3 | 6 | 2 | 3 | 1 | 5 | 49 | |
| V4a2 | _2007-16-25-0 | 40 | 3 | 15 | 2 | 40 | 4 | 4 | 241 | |
| | AB_2007-16-25- | 40 | 3 | 15 | 2 | 40 | 4 | 4 | 241 | |
| | AB_TG27339 | 40 | 3 | 15 | 2 | 40 | 4 | 4 | 241 | |
| V4a3 | UMB002 | 7 | 7 | 2 | 2 | 8 | 4 | 4 | 16 | |
| V4a4 | NIPH 60 | 8 | 1 | 14 | 3 | 12 | 1 | 13 | 34 | |
| V4a5 | NIPH 201 | 3 | 2 | 15 | 6 | 6 | 4 | 5 | 38 | |
| v4_1a1 | DSM30011 | 3 | 3 | 105 | 6 | 4 | 2 | 5 | 738 | |
| **V5 group** | | | | | | | | | | |
| V5a1 | BZICU-2 | 1 | 5 | 40 | 2 | 7 | 1 | 1 | 218 | |
| | Naval72 | 5 | 3 | 16 | 4 | 29 | 1 | 60 | 405 | |
| | TG22198 | nd | nd | nd | nd | nd | nd | nd | | |

[a]ST profiles (Pasteur Scheme) were retrieved from https://pubmlst.org/bigsdb?db=pubmlst_abaumannii_isolates&page=query

[b]Pertenence to the indicated global clonal complex (CC) as defined in ref. 8. A CC is a set of STs that are supposed to descend from the same found

[c]L1 or L2: sublineage 1 or 2 among CC1 as determined in ref. 9.

na: not available in the databas
