## Supplementary material for "Microevolution in the major outer membrane protein OmpA of *Acinetobacter baumannii*": Table S2

**Table S2. Nucleotide and amino acid mutations that distinguish the different *Acinetobacter ompA* variant alleles**

| Variant/allele | Length (nt/AA) | Number of Acinetobacter sp. strains with this gene allele | Number of A. baumannii strains with this gene allele | % of total A. baumannii strains (221) | % of A. baumannii V1 strains (143) | Nt changes as compared to V1a1 ompA | % identity to V1a1 | Amino acid residues changes as compared to V1a1 OmpA |
|---|---|---|---|---|---|---|---|---|
| V1 | 1,071/356 | 139 | 139 | 62.9 | 100 | | | |
| V1a1 | 1,071/356 | | 102 | 46.2 | 70.6 | none | 100.00 | none |
| V1a2 | 1,071/356 | | 1 | 0.45 | 0.70 | 1 (A564T) | 99.91 | none |
| V1a3 | 1,071/356 | | 3 | 1.36 | 2.10 | 7 (C396T, C414T, A431C, C432T, A435T, T489C, T747C) | 99.34 | 1 (T144N) |
| V1a4 | 1,071/356 | | 4 | 1.86 | 2.80 | 8 (T88C, C396T, C414T, A431C, C432T, A435T, T489C, T747C) | 99.25 | 1 (T144N) |
| V1a5 | 1,071/356 | | 2 | 0.90 | 1.40 | 8 (T88C, G154T, C186T, A210T, C396T, C414T, T747C, T903C) | 99.25 | 1 (A52S) |
| V1a6 | 1,071/356 | | 3 | 1.36 | 2.10 | 8 (T88C, C153T, G154T, G155C, T156A, C186T, C396T, C414T) | 99.25 | 1 (G52S) |
| V1a7 | 1,071/356 | | 2 | 0.90 | 1.40 | 8 (T88C, C153T, G154T, G155C, T156A, C186T, C396T, C414T, T747C) | 99.25 | 1 (G52S) |
| V1a8 | 1,071/356 | | 6 | 2.71 | 4.20 | 10 (T88C, C153T, G154T, G155C, T156A, C186T, C396T, C414T, T747C, T903C) | 99.07 | 1 (G52S) |
| V1a9 | 1,071/356 | | 3 | 1.36 | 2.10 | 11 (T88C, T144C, C153T, G154T, G155C, T156A, C186T, C396T, C414T, T747C, T903C) | 98.97 | 1 (G52S) |
| V1a10 | 1,071/356 | | 2 | 0.90 | 1.40 | 8 (T88C, G154T, C186T, A210T, T252C, C396T, A431C, A435T) | 99.25 | 2 (A52S, N144T) |
| V1a11 | 1,071/356 | | 1 | 0.45 | 0.70 | 9 (T88C, G154T, C198T, A210T, T252C, C396T, A431C, A435T, T747C) | 99.16 | 2 (A52S, N144T) |
| V1a12 | 1,071/356 | | 1 | 0.45 | 0.70 | 10 (T88C, G154T, C186T, A210T, T252C, C390T, G393A, A431C, C432T, A435T) | 99.07 | 2 (A52S, N144T) |
| V1a13 | 1,071/356 | | 3 | 1.36 | 2.10 | 13 (T88C, C153T, G154T, G155C, T156A, C186T, C396T, C414T, A431C, C432T, A435T, T747C, T903C) | 98.79 | 2 (G52S, N144T) |
| V1a14 | 1,071/356 | | 1 | 0.45 | 0.70 | 14 (T88C, C153T, G154T, G155C, T156A, C186T, C396T, C414T, A431C, C432T, A435T, T747C, C804T, T903C) | 98.69 | 2 (G52S, N144T) |
| V1a15 | 1,071/356 | | 1 | 0.45 | 0.70 | 10 (T88C, C153T, G154T, G155C, T156A, C186T, C396T, C414T, A646G, T747C) | 99.07 | 2 (G52S, T216A) |
| V1a16 | 1,071/356 | | 1 | 0.45 | 0.70 | 20 (T88C, C153T, G154T, A210T, G258T, C333T, A363G, G372A, C390T, G393A, C414T, A431C, C432T, A435T, T456C, G459T, T489C, A507T, C528T, T567C) | 98.13 | 3 (G52S, A90S, N144T ) |
| V1a17 | 1,071/356 | | 2 | 0.90 | 1.40 | 20 T88C, C153T, G154T, G155C, A210T, G268T, C333T, A363G, G372A, C390T, G393A, C414T, A431C, C432T, A435T, T456C, G459T, T489C, T747C, T903C) | 98.13 | 3 (G52S, A90S, N144T) |
| V1a18 | 1,071/356 | | 1 | 0.45 | 0.70 | 11 (T88C, G154T, C186T, A210T, T252C, C396T, A431C, A435T, T747C, T903C, A1051G) | 98.97 | 3 (A52S, N144T, T351A) |
| V1a19 | 1,071/356 | | 1 | 0.90 | 1.40 | 18 (T88C, C153T, G154T, G155C, T156A, C186T, C244T, G245C, T248G, C249T, C396T, T399C, C414T, A431C, C432T, A435T, T747C, T903C) | 98.32 | 4 (G52S, P89A, V90S, N144T) |
| | | | | % of A. baumannii V1 + V1_1 strains (149) | | | | |
| V1_1 | 1,053/350 | 6 | 4 | 1.81 | 2.68 | | | |
| V1_1a1 | 1,053/350 | | 1 | 0.45 | 0.67 | 29 (T88C, G154T, G155C, T156A, C186T, A210T, T252C, C396T, C414T, T894C, C1042G, 18 nt deletion at the C-terminal coding region) | 97.29 | 8 (A52S, Q348E, 6 aa deletion at the C-terminus) |
| V1_1a2 | 1,053/350 | | 3 | 1.36 | 2.01 | 28 (T88C, C153T, G154T, G155C, T156A, C186T, C396T, C414T, T894C, C1042G, 18 nt deletion at the C-terminal coding region | 97.39 | 8 (G52S, Q348E, 6 aa deletion in the C-terminus) |
| V1_1a3 | 1,053/350 | 1 (A. seifertii GG2) | | | | 77 (T88C, T123C, C145T, C153T, G154T, G155C, T156A, C198T, G199A, T201C, A204G, A210T, T252C, T255C, A259G, C285T, C317T, C333T, T345C, T355A, T357C, T366C, G375A, C390T, C396T, T408C, A431C, T456C, A463G, C465T, G471T, A476G, T477G, A491C, A523G, C525T, A531T, T516C, A536G, T537G, A541T, C543A, T546C, T549C, A567T, G578C, C588T, T589G, T590C, C598G, A600G, T627C, G774A, G822A, G825A, A829T, T897C, T903C, C1042G, deletion 18 nt at the C-terminal coding region) | 92.81 | 19 (G52S, I67V, N87D, A106V, F119I, N144T, I155V, Y159W, I161L, G173A, F177A, T257S, Q348E, 6 aa deletion at the C-terminus) |
| V1_1a4 | 1,053/350 | . npsocomialis OIFC021) | | | | 72 (T88C, T123C, C145T, C153T, G154T, G155C, T156A, G199A, T201C, A204G, A210T, T252C, T255C, A259G, C285T, C317T, C333T, T345C, T355A, T357C, T366C, G375A, C390T, C396T, T408C, A431C, C444T, T456C, A463G, C465T, A471T, A476G, T477G, A491C, C504T, T516C, A523G, C525T, G531T, A536G, T537G, A541T, C543A, T549C, A567T, G578C, C588T, T589G, T590C, A600G, T627C, T897C, T1005C, C1042G, deletion 18 nt at the C-terminal coding region) | 93.28 | 19 (G52S, I67V, N87D, A106V, F119I, N144T, I155V, Y159W, I161L, G173A, F177A, Q180E, Q348E, 6 aa deletion at the C-terminus) |
| | | | | | | Nt changes as compared to V2a1 ompA | % identity to | AA changes as compared to V2a1 OmpA |
| V2 | 1,062/353 | 58 | 57 | 25.8 | 100 | | | |
| V2a1 | 1,062/353 | | 38 | 17.2 | 66.7 | none | 100 | none |
| V2a2 | 1,062/353 | | 8 | 3.62 | 14.0 | 1 (T99C) | 99.91 | none |
| V2a3 | 1,062/353 | | 2 | 0.90 | 3.5 | 1 (T396C) | 99.91 | none |
| V2a4 | 1,062/353 | | 1 | 0.45 | 1.75 | 1 (A579T) | 99.91 | none |
| V2a5 | 1,062/353 | | 1 | 0.45 | 1.75 | 1 (C1002T) | 99.91 | none |
| V2a6 | 1,062/353 | | 2 | 0.90 | 3.5 | 1 (T268G) | 99.91 | 1 (S90A) |
| V2a7 | 1,059/352 | .calcoaceticus ANC3811) | | | | 27 (3 nt deletion at the transit peptide coding sequence (pos. 29-31 ), C39T, A45T, T48A, T75A, A81T, C163T, A259G, C264T, A288G, T408C, G412T, C417T, T454G, T484C, T558C, G753A, G756A, A760T, A825G, T894C, A897T, C963T, T1017A) | 97.45 | 5 (one deletion at the transit peptide, N87D, A138S, F152V, T253S) |
| | | | | % of A. baumannii V3 plus | | Nt changes as compared to V3a1 ompA | % identity to V3a1 ompA | |
| V3 | 1,029/342 | 17 | 9 | 4.1 | 100 | | | |
| V3a1 | 1,029/342 | | 9 | 4.1 | 100 | none | 100 | none |

| | | | | | | | | |
|---|---|---|---|---|---|---|---|---|
| V3a2 | 1,029/342 | lis WC-487, A. nosocomialis NIPH386) | | | 0 | 64 ( G36T, T65A, A90G, G91C, C99T, T120C, G127A, A138T, A139G, G140A, A153G, C168T, C169G, A180T, A182G, A192G, T198A, C228T, T234A, G237A, A261T, G262C, C263A, T264A, T267C, A270G, G274A, C306T, A315T, G321A, G330A, C336T, G354A, T354C, A363G, T372G, A390G, C399T, T402C, C407A, A414G, C432T, T435C, A439G, C441A, G447T, A459G, A471G, T483A, T492C, T498G, C504T, C510T, T525C, T528C, A531T, T535C, A537T, T543C, T558A, A708G, G810A, T873C, T981C) | 93.78 | 8 (V31L, D43N, S47D, A60G, A88Q, V97I, A136E, I147V) |
| V3a3 | 1,029/342 | , nosocomialis Ab22222) | | | 0 | 65 (G36T, T65A, A90G, G91C, C99T, T120C, G127A, A138T, A139G, G140A, A153G, C168T, C169G, A180T, A182G, A192G, T198A, C228T, T234A, G237A, A261T, G262C, C263A, T264A, T267C, A270G, G274A, C306T, A315T, G321A, G330A, C336T, G354A, T354C, A363G, T372G, A390G, C399T, T402C, C407A, A414G, C432T, T435C, A439G, C441A, G447T, A459G, A471G, T483A, T492C, T498G, C504T, C510T, T525C, T528C, A531T, T535C, A537T, T543C, T558A, A654T, A708G, G810A, T873C, T981C) | 93.68 | 8 (V31L, D43N, S47D, A60G, A88Q, V97I, A136E, I147V), OmpA protein identical to V3a2) |
| V3a4 | 1,026/341 | 7347, A. pittii 528, A. pittii TG2027) | | | 0 | 70 (3 nt deletion (pos. 29-31) at the transit peptide coding sequence, C39T, G66T, T75A, T84A, C88T, A90G, G91C, A93T, G108A, T111C, T114C, C117G, G127A, A138T, A139G, G140A, A153G, C168T, C179G, A180T, A249T, A261T, G262C, G263A, T264A, T267C, T268G, A273G, T276C, G279A, A282G, A285G, C306T, A315T, C318T, G330A, C336T, C348T, G354A, T357C, A363G, A372G, T441A, C455T, T458C, C465T, A468T, C469T, A471G, A473T, T492C, T498G, C510T, T525C, T528C, A531T, T543C, T558A, A708G, G738A, G741A, A745T, G810A, T879C, A882T, G936A, G999A | 93.20 | 9 ( 1 aa deletion at the transit peptide, V31L, D39E, D43N, S47D, A60G, A88Q, S90A, T249S) |
| V3a5 | 1,026/341 | . calcoaceticus RUH2202) | | | 0 | 71 (3 nt deletion (pos. 29-31) at the transit peptide coding sequence, C39T, A45T, T48A, T75A, C81T, T84A, G91C, A93T, G108A, T111C, T114C, T120C, G127A, A138T, A139G, A153G, G165A, C168T, C179G, A180T, A192G, T198A, C211T, C228T, T234A, A261T, G262C, G263A, T264A, T267C, A273G, T276C, A282G, A285G, C306T, T309A, A315T, G321A, T324C, G330A, C336T, C348T, G354A, A363G, A372G, C407A, A423T, T435C, T456C, A468T, C469T, A471G, A474T, T483A, T492C, T498G, T525C, T528C, A531T, T543C, T558A, A585T, T693A, T696C, G741A, A745T, T979C, A982T) | 93.10 | 8 (1 aa deletion at the transit peptide, V31L, D43N, S47D, A60G, A88Q, A136E, T249S) |
| V3a6 | 1,026/341 | A. calcoaceticus NIPH13) | | | 0 | 68 (3 nt deletion (pos. 29-31) at the transit peptide coding sequence, C39T, A45T, T48A, T75A, C81T, T84A, G91C, A93T, G108A, T111C, T114C, T120C, G127A, A138T, A153G, G165A, C168T, C179G, A180T, A192G, T198A, C211T, C228T, T234A, A261T, G262C, C263A, T264A, T267C, A273G, T276C, A282G, A285G, C306T, T309A, A315T, G321A, T324C, G330A, C336T, C348T, G354A, A363G, A372G, C407A, A423T, T435C, T456C, A468T, A471G, A474T, T483A, T492C, T498G, T525C, T528C, A531T, T543C,T558A, T693A, T696C, G741A, A745T, T979C, A982T) | 93.39 | 7 (1 aa deletion at the transit peptide, V31L, D43N, A60G, A88Q, A136E, T249S) |
| V3_1 | 1,044/347 | 7 | 0 | 0 | 0 | | | |
| V3_1a1 | 1,044/347 | 1 (A. pittii PHEA-2) | | | | 95 (3 nt deletion (pos. 29-31), C39T, G66T, T75A, T84A, C88T, A90G, G91C, A93T, G108A, T111C, T114C, C119G, G127A, A138T, A139G, G140A, A155G, C148T, C179G, A180T, A183G , A192G, T198A, C210T, C228T, T234A, C240T, A249T, A261T, G262C, C263A, T264A, T267C, T268G, A273C, T276G, G279A, A282G, A285G, C306T, A315T, C318T, G321A, C336T, C348T, G354A, T357C, A363G, A372G, C420T, T441A, C453T, T456C, A468T, A474T, T492C, T498G, C510T, T525C, T528C, A531T, T543C, T558A, A615T, A708G, G738A, G741A, A745T, G810A, T879C, A882T, T1002A, G1118C, G1023T, 18 nt (GCAGCTCCTGCAGCAGCT) insertion at the C-terminal coding region) | 90.90 | 16 ( 1 aa deletion at the transit peptide, V31L, D39E, S47D, A60G, E83D, A88Q, S90A, T249S, E280Q , 6 aa insertion (AAAPAA) at the C-terminal region) |
| V3_1a2 | 1,044/347 | 1 (A. pittii 4050) | | | | 88 (3 nt deletion (pos. 29-31), C39T, G66T, T75A, T84A, A90G, G91C, A93T, G108A, T111C, T114C, C119G, G127A, A138T, A139G, G140A, A155G, C148T, C179G, A180T, A183G, A192G, T198A, C213T C228T, T234A, A249T, A261T, G262C, C263A, T264A, T267C, T268G, A273C, T276G, G279A, A282G, A285G, C306T, G321A, C336T, C348T, G354A, T357C, A363G, A372G, T456C, T462C, , A468T, A474T, T492C, T498G, C510T, T525C, A531T, T543C, T558A, T651C, A708G, G738A, G741A, A745T, G810A, T879C, A882T, T1002A, G1118C, G1023T, 18 nt (GCAGCTCCTGCAGCAGCT) insertion at the C-terminal coding region) | 91.57 | 16 ( 1 aa deletion at the transit peptide, V31L, D39E, S47D, A60G, E83D, A88Q, S90A, T249S, E280Q , 6 aa insertion (AAAPAA) at the C-terminal region), identical to V3_1_a1 |
| V3_1a3 | 1,044/347 | 1 (A. pittii 4052) | | | | 88 (3 nt deletion (pos. 29-31), C39T, G66T, T75A, T84A, A90G, G91C, G108A, T111C, T114C, C119G, G127A, A138T, A155G, C148T, C179G, A180T, A183G, A192G, T198A, C213T, C228T, T234A, A249T, A261T, G262C, C263A, T264A, T267C, T268G, A273C, T276G, G279A, A282G, A285G, C306T, A315T, G321A, C336T, C348T, G354A, T357C, A363G, A372G, C420T, T456C, A468T, A474T, T492C, T498G, C510T, T525C, T528C, A531T, T543C, T558A, T651C, A672G, A708G, G738A, G741A, A745T, G810A, T879C, A882T, T1002A, G1118C, A1023T, 18 nt (GCAGCTCCTGCAGCAGCT) insertion at the C-terminal coding region) | 91.57 | 15 ( 1 aa deletion at the transit peptide, V31L, D39E, A60G, E83D, A88Q, S90A, T249S, E280Q , 6 aa insertion (AAAPAA) at the C-terminal region) |

| | | | | | | | | |
|---|---|---|---|---|---|---|---|---|
| V3_1a4 | 1,044/347 | .calcoaceticus TG19593) | | | | 89 (3 nt deletion (pos. 29-31), C39T, A45T, T48A, T75A, C81T, T84A, G91C, A93T, G108A, T111C, T114C, C119G, G127A, A139G, G140A, A155G, C148T, C179G, A180T, A183G, A192G, T198A C213T, C228T, T234A, A261T, G262C, C263A, T264A, T267C, A273C, T276G, A282G, A285G, T300C, C306T, A315T, G321A, C336T, C348T, G354A, A363G, A372G, A423T, T435C, T456C, T462C, A468T, A474T, T483A, T492C, T498G, G513A, T525C, T528C, A531T, T543C, T558A, T696C, A708G, G738A, G741A, A745T, T879C, A882T, C948T, T1002A, G1118C, G1023T, 18 nt (GCAGCTCCTGCAGCAGCT) insertion at the C-terminal coding region) | 91.48 | 14 ( 1 aa deletion at the transit peptide, V31L, D39E, S47D, A60G, A88Q, T249S, E280Q, 6 aa insertion (AAAPAA) at the C-terminal region) |
| V3_1a5 | 1,044/347 | . calcoaceticus ANC 3680) | | | | 91 (3 nt deletion (pos. 29-31), C39T, A45T, T48A, T75A, C81T, T84A, G91C, A93T, G108A, T111C, T114C, C119G, G127A, A139G, G140A, A155G, C148T, C179G, A180T, A183G, A192G, T198A C213T, C228T, T234A, A261T, G262C, C263A, T264A, T267C, T268G, A273C, T276G, A282G, A285G, C306T, T309A, A315T, G321A, C336T, C348T, G354A, A363G, A372G, A423T, T435C, T456C, T462C, A468T, A474T, T483A, T492C, T498G, G513A, T525C, T528C, A531T, T543C, T558A, A708G, G738A, G741A, A745T, T879C, A882T, C948T, T1002A, G1118C, G1023T, 18 nt (GCAGCTCCTGCAGCAGCT) insertion at the C-terminal coding region) | 91.28 | 14 ( 1 aa deletion at the transit peptide, V31L, D39E, S47D, A60G, A88Q, T249S, E280Q, 6 aa insertion (AAAPAA) at the C-terminal region), identical to V3_1_a4 |
| V3_1a6 | 1,044/347 | 2 (A. oleivorans DR1, A. oleivorans CIP 110421) | | | | (3 nt deletion (pos. 29-31), C39T, A45T, T48A, T75A, C81T, T84A, G91C, A93T, G108A, T111C, T114C, C119G, G127A, A139G, G140A, A155G, C148T, C179G, A180T, A183G, A192G, T198A C213T, C228T, T234A, A261T, G262C, C263A, T264A, T267C, T268G, A273G, T276C, A282G, A285G, G289A, C306T, A315T, G321A, C336T, G354A, A363G, T372G, A423T, T435C, T456C, T462C, C465T, A468T, A474T, T483A, T492C, T498G, G513A, T525C, T528C, A531T, T543C, T558A, A585T, G690A, A708G, G738A, G741A, A745T, T879C, A882T, T1002A, G1118C, G1023T, 18 nt 91 (GCAGCTCCTGCAGCAGCT) insertion at the C-terminal coding region) | 91.28 | 14 ( 1 aa deletion at the transit peptide, V31L, D39E, S47D, A60G, A88Q, T249S, E280Q, 6 aa insertion (AAAPAA) at the C-terminal region), identical to V3_1_a4 |
| | | | | % of A. baumannii V4 plus | | Nt changes as compared to V4a1 ompA | | |
| V4 | 1,050/349 | 8 | 8 | 4.1 | 100 | | | |
| V4a1 | 1,050/349 | | 2 | 0.90 | 22 | none | | |
| V4a2 | 1,049/349 | | 3 | 1.36 | 33.3 | 1 (G270A) | 99.90 | Identical to V4a1 |
| V4a3 | 1,040/349 | | 1 | 0.45 | 11.1 | 10 (T249C, C250A, A252G, C258T, G270A, T285C, C465T, G675A, C726T, T909C) | 99.05 | 1 (P84T) |
| V4a4 | 1,038/349 | | 1 | 0.45 | 11.1 | 12 (T249C, C250A, A252G, C258T, G270A, T285C, G450T, C465T, C472T, T513C, G675A, C684T) | 98.86 | 1 (P84T) |
| V4a5 | 1,036/349 | | 1 | 0.45 | 11.1 | 14 ( T249C, C250A, A252G, C258T, G270A, T285C, C465T, C472T, A537T, G675A, C726T, T786C, T873C, C1021G) | 98.67 | 2 (P84T, Q341E) |
| V4_1 | 1,032/343 | 1 | 1 | | | | | |
| V4_1a1 | 1,032/343 | | 1 | 0.45 | 11.1 | 32 (T249C, C250A, A252G, C258T, G270A, T285C, A442G, C465T, A537T, G675A, C726T, T786C, T877C, C1021G, 18 nt deletion at the C-terminal coding region) | 95.24 | 9 (P84T, I148V, Q341E, 6 aa deletion (AAAPAA) at the C-terminal region) |
