## Supplementary material for "Microevolution in the major outer membrane protein OmpA of *Acinetobacter baumannii*": Table S3

**Table** S3. Identities and similarities between the different *A. baumannii* OmpA variants detected in this work. The numbers above and below the vertical line defined by complete (100 %) sequence identity indicate the percentages of identity and similarity, respectively, for each given pair at the amino acid sequence level.

| | ACICU (V1_a1) | TG02017 (V1_1_a1) | AB0057 (V2_a1) | 4190 (V3_a1) | AB900 (V4_a1) | DSM30011 (V4_1_a1) | BZICU-2 (V5_a1) |
|---|---|---|---|---|---|---|---|
| ACICU (V1_a1) | 100 | 97.5 | 94.1 | 84.6 | 92.4 | 89.9 | 91.9 |
| TG02017 (V1_1_a1) | 97.8 | 100 | 91.9 | 86.6 | 89.9 | 91.7 | 94.0 |
| AB0057 (V2_a1) | 95.8 | 94.1 | 100 | 85.1 | 92.4 | 89.9 | 92.1 |
| 4190 (V3_a1) | 89.4 | 91.2 | 89.6 | 100 | 86.9 | 88.7 | 86.7 |
| AB900 (V4_a1) | 94.4 | 92.1 | 94.1 | 90.9 | 100 | 97.1 | 94.9 |
| DSM30011 (V4_1_a1) | 92.4 | 94.0 | 92.1 | 92.8 | 97.7 | 100 | 96.5 |
| BZICU-2 (V5_a1) | 93.8 | 96.0 | 93.5 | 91.1 | 95.8 | 97.4 | 100 |
