## Supplementary material for "Microevolution in the major outer membrane protein OmpA of *Acinetobacter baumannii*": Table S4

**Table S4. N-terminal transit peptides and C-terminal sequences of OmpA proteins of *Acinetobacter* genus species.** The *Acinetobacter* species and the accession numbers of the corresponding OmpA proteins as well as the different ecologically-differentiated clades in which the genus was recently divided (ref. 1) are indicated in the first column. In the case of ACB complex members, the corresponding OmpA variant alleles (as determined in this work) are also indicated. The aligned sequences at the left show the amino acid sequences of the transit peptides in each case, with deletion/insertions indicated by hyphens (-) and the "▼" symbol the processing site for Signal peptidase I as inferred using http://www.cbs.dtu.dk/services/SignalP/). In turn, the aligned sequences at the right show the last amino acids of the C-terminal ends for the corresponding OmpAs.

| Species and clades | Transit peptide | C-terminus |
|---|---|---|
| A.baylyi ADP1 CAG67610.1 | MKLSRIALATMLVAAPLAAANA▼...... | TVTVQPGQQAPAAQ |
| A.ursingii WP_004991206.1 | MKLSRIALATMLVAAPFAAANA▼...... | TVVVQPGQQAQ |
| A.soli WP_076032993.1 | MKLSRIALATMLVAAPLAAANA▼...... | TVTVQPGQQAQ |
| **Clade I (ACB complex)** | | |
| **A. baumannii** | | |
| A.baumannii V1 alleles | MKLSRIALATMLVAAPLAAANA▼...... | TVVVQPGQEAAAPAAAQ |
| A.baumannii V1_1 alleles | MKLSRIALATMLVAAPLAAANA▼...... | TVVVQPGQQAQ |
| A.baumannii V2 alleles | MKLSRIALATMLVAAPLAAANA▼...... | TVVVQPGQEAAAPAAAQ |
| A.baumannii V3 alleles | MKLSRIALATMLVAAPLAAANA▼...... | TVVVQPGQQAQ |
| A.baumannii V4 alleles | MKLSRIALATMLVAAPLAAANA▼...... | TVVVQPGQEAAAPAAAQ |
| A.baumannii V4_1 alleles | MKLSRIALATMLVAAPLAAANA▼...... | TVVVQPGQQAQ |
| A.baumannii V5 alleles | MKLSRIALATMLVAAPLAAANA▼...... | TVVVQPGQQAQ |
| **Non-A.baumannii species** | | |
| A.nosocomialis OIFC021 (V1_1a4) | MKLSRIALATMLVAAPLAAANA▼...... | TVVVQPGQQAQ |
| A.nosocomialis WC-487 (V3a2) | MKLSRIALATMLVAAPLAAANA▼...... | TVVVQPGQQAQ |
| A.seifertii GG2 (V1_1a3) | MKLSRIALATMLVAAPLAAANA▼...... | TVVVQPGQQAQ |
| A.calcoaceticus ANC3811 (V2a7) | MKLSRIALA-MLVAAPLAAANA▼...... | TVVVQPGQEAAAPAAAQ |
| A.calcoaceticus RUH2202(V3a5) | MKLSRIALA-MLVAAPLAAANA▼...... | TVVVQPGQQAQ |
| A.calcoaceticus NIPH13 (V3a6) | MKLSRIALA-MLVAAPLAAANA▼...... | TVVVQPGQQAQ |
| A.calcoaceticus TG19593(V3_1a4) | MKLSRIALA-MLVAAPLAAANA▼...... | TVVVQPGQEAAAPAAAQ |
| A.oleivorans DR1 (V3_1a6) | MKLSRIALA-MLVAAPLAAANA▼...... | TVVVQPGQEAAAPAAAQ |
| A.pittii PHEA-2(V3_1a1) | MKLSRIALA-MLVAAPLAAANA▼...... | TVVVQPGQEAAAPAAAQ |
| A.pittii ANC4050 (V3_1a2) | MKLSRIALA-MLVAAPLAAANA▼...... | TVVVQPGQEAAAPAAAQ |
| A.pittii ANC4052 (V3_1a3) | MKLSRIALA-MLVAAPLAAANA▼...... | TVVVQPGQEAAAPAAAQ |
| A.pittii WC-136 (V5a2) | MKLSRIALA-MLVAAPLAAANA▼...... | TVVVQPGQQAQ |
| **Clade II** | | |
| A.junii WP_039047009.1 | MKLSRIALA-MLVAAPLAAANA▼...... | TVLVQPGQQAQ |
| A.beijerinckii WP_005061574.1 | MKLSRIALA-MLVAAPLAAANA▼...... | TVLVQPGQQAQ |
| A.parvus WP_050041456.1 | MKLSRIALA-MLVAAPLAAANA▼...... | TVLVQPGQQAQ |
| A.proteolyticus WP_101235588.1 | MKLSRIALA-MLVAAPLAAANA▼...... | TVLVQPGQQAQ |
| A.haemolyticus SPT46410.1 | MKLSRIALA-MLVAAPFAAANA▼...... | TVLVQPDQQAQ |
| A.venetianus WP_019383768.1 | MKLSRIALA-MLVAAPLAAANA▼...... | TVLVQPGQ |
| A.tjernbergiae WP_018677830.1 | MKLSRIALA-MLVAAPLAAANA▼...... | TVLVQPGQ |
| **Clade III** | | |
| A.gerneri WP_004854794.1 | MKLSRIALA-MLVAAPLAAANA▼...... | TVVQDAQ |
| A.bereziniae WP_004829936.1 | MKLSRIALA-MLVAAPLAAANA▼...... | TVTKTVTK |
| A.rudis EPF79807.1 | MKLSRIALA-MLVAAPLAAANA▼...... | TVLAQPRAQPR |
| A.indicus AQU14364.1 | MKMSRIALA-MLVAAPLAAANA▼...... | TVTVQPEAAAQ |
| A.schindleri WP_076754170.1 | MKMSRIALA-MLVAAPLAAANA▼...... | TVLAEQPAAQ |

```
A.lwoffii AUC08284.1           MKMSRIALA-MLVAAPFAAANA▼……TVLAEQPAQ
A.bouvetii WP_005007826.1      MKMSRIALA-MLVAAPLAAANA▼……TVVVEGQQAQ
A.johnsonii WP_058870218.1     MKMSRIALA-MLVAAPLAAANA▼……TVQAQ
A.nectaris WP_023273734.1      MKLSRIALATVLAASPFVVANA▼……TVLAQPKAQ
A.brisouii WP_045794742.1      MKLSRIAVATLLAASPLVAANA▼……TVIAQPTAPAAQ
```
