## Supplementary material for "Microevolution in the major outer membrane protein OmpA of *Acinetobacter baumannii*": Table S5

Table S5: Variation in substitution patterns between extracellular loop regions and non-exposed regions of *ompA* for total alignments (Recombination present) and alignments where regions identified as recombinant using RDP have been removed (Recombination removed). [a]Values for non-exposed protein domains ($\kappa1$ and $\omega1$) and exposed extracellular loops ($\kappa2$ and $\omega2$) are shown. $\kappa$, transition/transversion rate ratio; $\omega$, $d_N/d_S$ ratio; LRT, likelihood ratio test. [b]Significance assessed using $\chi^2$ tests, with *p* value corrected for multiple testing using a Bonferroni correction.

| ***ompA* clades** | **Fixed parameter model** | | **Variable parameter model[a]** | | **LRT statistic** | ***p* value[b]** |
|---|---|---|---|---|---|---|
| | $\kappa$ | $\omega$ | $\kappa$ | $\omega$ | | |
| *Recombination present* | | | | | | |
| V1 | 2.38998 | 0.05882 | $\kappa1$: 4.10329 | $\omega1$: 0.04248 | 22.309152 | 0.00004293 |
| | | | $\kappa2$: 1.10660 | $\omega2$: 0.06264 | | |
| V2 | 5.01699 | 0.03137 | $\kappa1$: 10.60187 | $\omega1$: 0.00010 | 28.942516 | 0.000001557 |
| | | | $\kappa2$: 0.82641 | $\omega2$: 0.17593 | | |
| V4 | 4.39979 | 0.04919 | $\kappa1$: 4.87079 | $\omega1$: 0.05020 | 22.975272 | 0.000030768 |
| | | | $\kappa2$: 2.57351 | $\omega2$: 0.04172 | | |
| *Recombination removed* | | | | | | |
| V1 | 3.05724 | 0.12321 | $\kappa1$: 1.85014 | $\omega1$: 0.10181 | 3.839502 | 0.587 |
| | | | $\kappa2$: 6.41894 | $\omega2$: 0.14789 | | |
| V2 | 5.33917 | 0.03150 | $\kappa1$: 5.65768 | $\omega1$: 0.03506 | 12.915742 | 0.006 |
| | | | $\kappa2$: 999.00000 | $\omega2$: 0.00010 | | |
| V4 | 4.27961 | 0.04928 | $\kappa1$: 4.12621 | $\omega1$: 0.05526 | 7.48023 | 0.095 |
| | | | $\kappa2$: 4.72598 | $\omega2$: 0.04453 | | |
