## Supplementary material for "Microevolution in the major outer membrane protein OmpA of *Acinetobacter baumannii*": Table S6

Table S6: Bacteria with close matches to the 18 nucleotide high-GC sequence coding for an alanine-rich tract located at the c-terminal end of some OmpA variants.

| Phyla | Species | GenBank Accession number |
|---|---|---|
| *Actinobacteria* | *Streptomyces* sp. HNM0039 | CP029188 |
| | *Propionibacterium australiense* | LR134442 |
| | *Gordonia* spp. | CP022580, CP002907 |
| | *Pseudonocardia* spp. | CP010989, CP012181, CP012184 |
| | *Stackebrandtia nassauensis* | CP001778 |
| *Bacteroidetes* | *Hymenobacter* APR13 | CP006587 |
| *Proteobacteria* | *Pseudomonas stutzeri* | CP007441 |
| | *Serratia ficaria* | LT906479 |
| | *Comamonas* spp. | LN879547, CP001220 |
| | *Bradyrhizobium canariense* | LT629750 |
| | *Martelella endophytica* | CP010803 |
| | *Novosphingobium* THN1 | CP028347 |
